# Ultra-high efficiency T cell reprogramming at multiple loci with SEED-Selection

**DOI:** 10.1101/2024.02.06.576175

**Authors:** Christopher R. Chang, Vivasvan S. Vykunta, Daniel B. Goodman, Joseph J. Muldoon, William A. Nyberg, Chang Liu, Vincent Allain, Allison Rothrock, Charlotte H. Wang, Alexander Marson, Brian R. Shy, Justin Eyquem

## Abstract

Multiplexed reprogramming of T cell specificity and function can generate powerful next-generation cellular therapies. However, current manufacturing methods produce heterogenous mixtures of partially engineered cells. Here, we develop a one-step process to enrich for unlabeled cells with knock-ins at multiple target loci using a family of repair templates named <u>S</u>ynthetic <u>E</u>xon/<u>E</u>xpression Disruptors (SEEDs). SEED engineering associates transgene integration with the disruption of a paired endogenous surface protein, allowing non-modified and partially edited cells to be immunomagnetically depleted (SEED-Selection). We design SEEDs to fully reprogram three critical loci encoding T cell specificity, co-receptor expression, and MHC expression, with up to 98% purity after selection for individual modifications and up to 90% purity for six simultaneous edits (three knock-ins and three knockouts). These methods are simple, compatible with existing clinical manufacturing workflows, and can be readily adapted to other loci to facilitate production of complex gene-edited cell therapies.

## Main

T cells engineered to express synthetic immune receptors are highly effective for the treatment of refractory hematological malignancies^1,2^. Nevertheless, efforts to create new cell therapies have been stymied by difficulties maintaining T cell persistence and functionality^3,4^. Furthermore, widespread adoption of cell therapies has been hindered by the autologous nature of current products, which require an expensive and time-consuming individualized manufacturing process^5^.

Combinations of transgenes integrated into the genome via CRISPR/Cas editing have been used to improve the performance of cell therapies or to create allogeneic (off-the-shelf) products^6–9^. However, viral and non-viral DNA repair templates have a limited cargo capacity, which constrains the number and size of transgenes that can be introduced at one locus^10,11^. CRISPR/Cas can be used to introduce targeted double-strand breaks (DSBs) at multiple loci simultaneously, but achieving multiple transgene integrations is challenging since non-homologous end joining (NHEJ) (which creates insertions and deletions) generally outcompetes homology-directed repair (HDR) (which facilitates transgene integration)^12^. As a result, efforts to perform multiplexed knock-ins have yielded numerous populations of partially-edited cells that can perform sub-optimally^13–16^. Since product purity is critical for clinical manufacturing, efficient methods for isolating fully edited cells are necessary for the realization of multi-locus integration strategies.

Isolating engineered cells has been a long-lasting interest of the field, and a variety of methods have been developed. Surface tags and drug resistance cassettes have been used to enrich for cells with transgene integrations, but subjecting cells to multiple drugs or performing sequential rounds of positive selection can negatively impact cell viability, performance, and yield^16–19^. Alternatively, the targeting of essential loci has been used to enrich for cells with transgene integrations^20^, but the consequences of simultaneously editing multiple essential genes have not yet been evaluated. As editing outcomes at distinct loci are linked, previous studies have used a selective marker introduced at one locus to enrich for integrations at another locus^15,21,22^. However, an enrichment method for direct multi-marker selection would allow for the isolation of purer populations.

Here, we develop a one-step, drug-free process to isolate unlabeled cells that have transgene integrations at multiple loci. We devise a type of repair template named a <u>S</u>ynthetic <u>E</u>xon/<u>E</u>xpression Disruptor (SEED) to link successful transgene integration with the disruption of a paired endogenous surface protein, allowing cells with knock-ins to be enriched through immunomagnetic negative selection (SEED-Selection).

We design SEEDs to disrupt three translationally relevant surface proteins in primary human T cells while facilitating the expression of various therapeutic payloads. We characterize editing outcomes and transgene function in cells edited with a single or multiple SEEDs, and the ability of SEED-Selection to enrich for cells with biallelic integrations in a single step. Additionally, we demonstrate that antibody epitope editing enables the enrichment of transgenes that would otherwise be depleted during SEED-Selection and facilitates the removal of T cells with mispaired T cell receptors (TCRs) through SEED-Selection when a transgenic TCR is introduced at the TCR α constant (*TRAC*) locus.

SEED-Selection facilitates the isolation of almost entirely pure (up to 98%) populations of cells with an intended knock-in and knockout. Furthermore, SEED-selection is amenable to multiplexing, allowing for the isolation of highly pure (up to 90% fully edited) populations that have three knock-ins and three knockouts. SEED-selection could be easily adapted to various cell types to facilitate clinical manufacturing for a wide variety of complex gene-edited cell therapies.

## Results

### SEED engineering enables efficient enrichment of cells with transgene integrations

To develop a method that would allow for cells with transgene integrations to be enriched through negative selection, we designed two reagents: (1) a guide RNA (gRNA) targeting an intron of a surface-expressed protein that generates a DSB that minimally impacts expression, and (2) a SEED homology-directed repair template (HDRT) that utilizes synthetic splice acceptor (SA) and splice donor (SD) sequences to introduce an in-frame transgene payload preceded by a P2A sequence at a position that disrupts the target protein (Fig. 1a).

**Figure 1:**
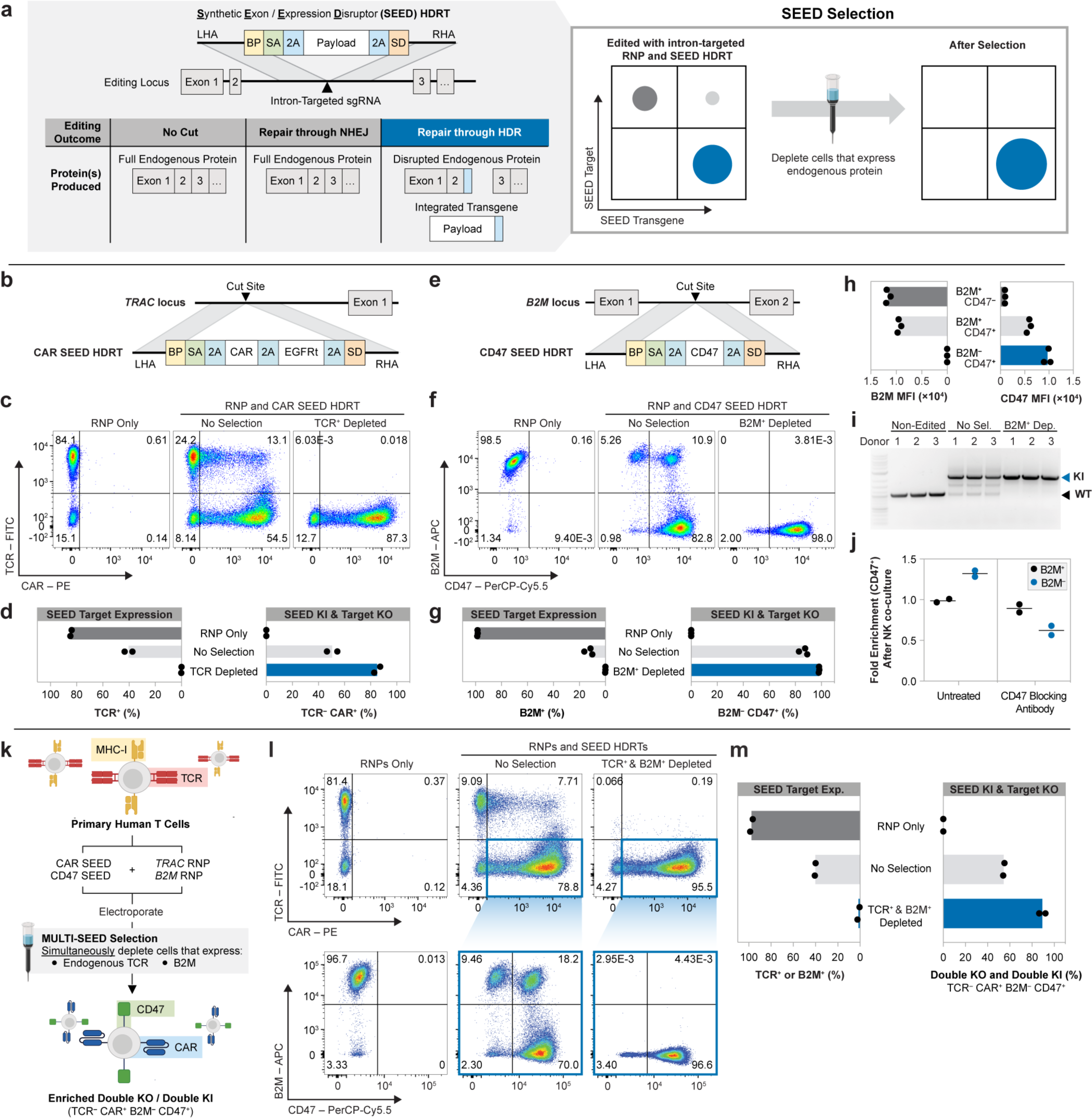
SEED-Selection enriches for cells with integrations at multiple loci. **a**, Overview of editing outcomes generated with an intron-targeted gRNA and a SEED HDRT. Immunomagnetic reagents are used to deplete cells that retain expression of the surface protein targeted by a SEED, thereby enriching for cells with transgene integration. **b**, Diagram of a *TRAC* intron-targeted SEED HDRT encoding a CAR and EGFRt. **c,d,** T cells were edited with *TRAC* intron-targeted RNP and HDRT (**b**) then immunomagnetically purified with anti-TCR (*n* = 2 donors). **c,** Flow cytometry plots of TCR and CAR expression (anti-G4S linker). **d,** Percentage of TCR^+^ and TCR^-^ CAR^+^ cells. **e**, Diagram of a *B2M* intron-targeted SEED HDRT encoding CD47. **f–i,** T cells were edited with *B2M* intron-targeted RNP and HDRT) then immunomagnetically purified with anti-B2M (*n* = 3 donors). **f**, Flow cytometry plots of B2M and CD47 expression. **g,** Percentage of B2M^+^ and B2M^-^ CD47^+^ cells. **h**, Expression of B2M or CD47 in subpopulations of non-purified edited cells. **i,** Genomic DNA PCR targeting the SEED integration site at *B2M*. Amplicon for non-edited alleles (black triangle); amplicon for HDRT integration (blue triangle). **j,** Fold enrichment of CD47^+^ SEED-edited cells relative to CD47^-^ cells after overnight co-culture with NK cells. **k–m,** T cells were edited with *TRAC* and *B2M* RNPs and transduced with *TRAC*-CAR SEED (**b**) and *B2M*-CD47 SEED (**e**) HDRTs. Edited cells were then immunomagnetically purified with anti-B2M and anti-TCR (*n* = 2 donors). **k**, Diagram of multiplexed editing and enrichment strategy. **l,** Representative flow cytometry plots of TCR, CAR, B2M, and CD47 expression. Colored boxes indicate subpopulations in each sample. **m,** Percentage of B2M^+^ or TCR^+^ cells and fully edited cells (TCR^-^ CAR^+^ B2M^-^ CD47^+^). BP: branchpoint; SA: splice acceptor; SD: splice donor; LHA/RHA: homology arms; MFI: median fluorescence intensity.

We initially targeted the two most common loci for therapeutic T cell engineering, *TRAC* and *B2M*. Insertion of a chimeric antigen receptor (CAR) or transgenic TCR at the *TRAC* locus can directly reprogram T cell specificity, benefitting from endogenous regulatory elements to generate potent and durable T cell therapies^23,24^. B2M is required for MHC-I expression and can be disrupted to help evade host T cell recognition in allogeneic settings^25^. To identify optimal SEED integration sites for *TRAC* and *B2M*, we designed two panels of intron-targeted gRNAs and screened for guides with the ability to generate insertions and deletions (indels) in primary human T cells without disrupting TCR or B2M surface expression, respectively (Extended Data Fig. 1a–d). Candidate intronic gRNAs exhibited high indel generation efficiencies (up to 97%) while minimally disrupting TCR (9–16%) or B2M (1–2%) surface expression.

To confirm that SEED integration could disrupt TCR surface expression, we edited T cells with a *TRAC* intron-targeted RNP and an adeno-associated viral vector (AAV6) designed to deliver a SEED HDRT encoding a CD19-specific CAR and a truncated EGFR (EGFRt) surface receptor (Fig. 1b,c)^18^. Editing was performed in the presence of a DNA-PK inhibitor (M3814), which has been shown to improve HDR efficiency by blocking NHEJ^11,26^. As intended, TCR disruption in cells edited with the RNP alone was infrequent (16%) (Fig. 1d). Integration-mediated TCR disruption (TCR^-^ CAR^+^) occurred in half (50.5%) of SEED-transduced cells, which were enriched to purities >85% through immunomagnetic TCR depletion (Fig. 1d).

To compare editing outcomes for SEED and exon-targeting strategies, we designed and tested an analogous CAR HDRT with an integration site in *TRAC* exon 1 (Extended Data Fig. 2a,b)^18^. CAR expression levels and homogeneity were similar in T cells edited with both constructs (Extended Data Fig. 2c,d). However, immunomagnetic TCR depletion of cells edited with an exon-targeted HDRT minimally enriched for TCR^-^ CAR^+^ cells, since the majority of CAR^-^ cells were also TCR^-^ (Extended Data Fig. 2e–g). These data demonstrate that SEEDs enable edited cells to be enriched by coupling transgene integration to target protein disruption.

### SEED-Selection enriches for cells with biallelic integrations

As the *TRAC* locus is subject to allelic exclusion, editing at the single active allele is often sufficient to ablate TCR expression^23^. In contrast, editing at both alleles is necessary to fully disrupt B2M expression. Therefore, we hypothesized that SEED engineering and selection strategies targeting B2M would enrich for cells with biallelic SEED integrations.

We designed an AAV6 SEED HDRT to simultaneously disrupt B2M expression and deliver CD47 – an immune checkpoint molecule that has previously been shown to inhibit NK and macrophage activity against cells which lack MHC-I expression (Fig. 1e)^27–29^. Remarkably, integration-mediated B2M disruption (B2M^-^ CD47^+^) was achieved in >85% of cells edited with a *B2M* intron-targeted RNP and SEED HDRT, while minimal B2M disruption (<2%) occurred when editing was performed with the RNP alone (Fig. 1f,g). Although most cells that overexpressed CD47 had complete disruption of B2M, a small subpopulation expressed intermediate levels of B2M and CD47 (Fig. 1h), consistent with monoallelic SEED integration. Immunomagnetic selection efficiently removed cells with endogenous and intermediate levels of B2M expression, allowing for the isolation of a pure (>98%) B2M^-^ CD47^+^ population with biallelic SEED integration (Fig. 1f,g). Product purity was further evaluated through genomic DNA PCR, which confirmed depletion of non-edited *B2M* alleles during selection (Fig. 1i).

To test whether CD47 expression in SEED-edited T cells was sufficient to reduce NK cell cytotoxicity, we generated a mixture of T cells with four subsets (B2M^+^ CD47^-^; B2M^+^ CD47^+^; B2M^-^ CD47^+^; B2M^-^ CD47^-^) and performed a co-culture with activated primary human NK cells. The composition of the co-culture was then quantified via flow cytometry and compared to T cells alone (Extended Data Fig. 3). As intended, B2M^-^ CD47^+^ cells were enriched relative to B2M^-^ CD47^-^ cells after co-culture (Fig. 1j). B2M^-^ CD47^+^ enrichment was not observed when T cells were pre-treated with a CD47-blocking antibody, confirming that SEED-mediated CD47 activity facilitated NK cell evasion (Fig. 1j).

### SEED-Selection can simultaneously enrich for edits at multiple loci

Multiplexed immunomagnetic selection kits are commonly used to isolate cell subsets, and customized selection panels can be created by mixing antibodies targeting markers of interest. To test whether SEEDs targeting different loci could be enriched in a single step, we performed multiplexed editing to introduce *TRAC*-CAR and *B2M*-CD47 SEEDs into T cells (Fig. 1k). High integration efficiencies were observed at both loci, with up to 55% of cells undergoing full editing (TCR^-^ CAR^+^ B2M^-^ CD47^+^) (Fig. 1l,m). Simultaneous immunomagnetic purification with TCR and B2M-targeted antibodies further enriched for cells with transgene integrations, allowing for the isolation of highly pure (up to 92%) cells with two knockouts and two knock-ins (Fig. 1m and Extended Data Fig. 4). These data demonstrate that SEED-Selection can be multiplexed without compromising product purity.

### Epitope editing allows for TCR-based receptors to evade TCR-targeted antibodies

Synthetic immune receptors containing TCRα/β constant domains, such as transgenic α/β TCRs^30^ or HLA-independent TCRs (HITs)^31^, are highly sensitive to low antigen densities. However, these receptors are incompatible with TCR SEED-Selection since they are also bound by the α/β TCR-targeted antibodies used to deplete non-modified and partially edited cells (Fig. 2a–c). Structural analysis and high-throughput screening techniques have been used to design epitope-edited receptors that evade a specific antibody and remain functional^32–34^. We hypothesized that epitope editing of SEED payloads would allow for the enrichment of cells with a transgene that would otherwise be depleted through SEED-Selection.

**Figure 2:**
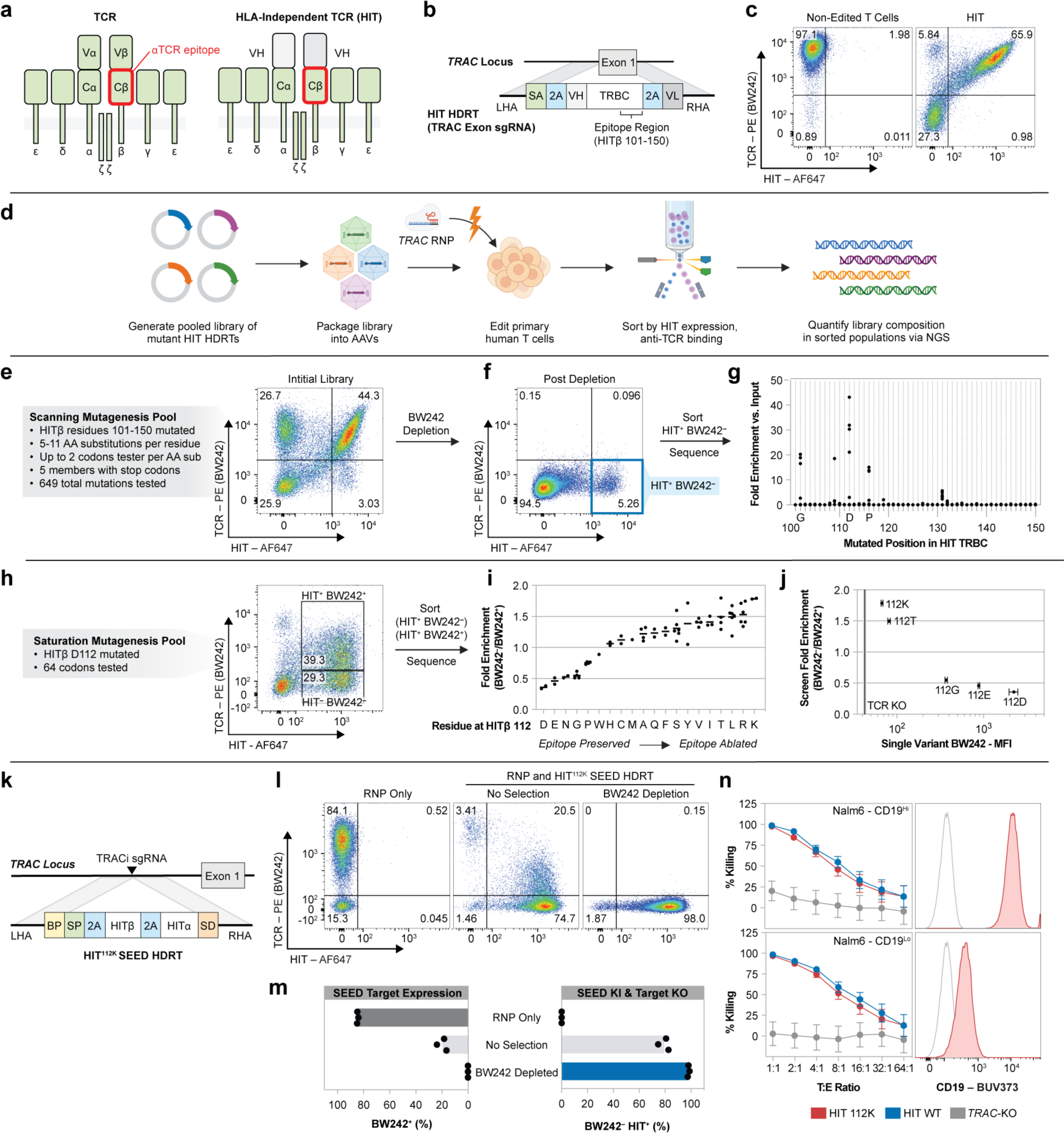
Epitope editing allows for TCR-based receptors to be used in TCR SEEDs. **a**, Diagram of a TCR and a HIT. **b**, Diagram of a *TRAC* exon-targeted HDRT encoding a HIT. **c**, Flow cytometry plot of anti-TCR (BW242) binding and HIT expression (anti-mouse F(ab’)_2_) in T cells with a HIT introduced at *TRAC* (*n* = 1 donor). **d**, Pooled screening workflow for epitope mapping. **e–g**, T cells were edited to express a library of mutated HITs then immunomagnetically purified with BW242. Flow cytometry was used to quantify BW242 binding and HIT expression before (**e**) and after (**f**) purification. HIT^+^ BW242^-^ cells were sorted (blue box) for sequencing analysis. **g**, Relative enrichment of mutations in HIT^+^ BW242^-^ cells (blue box in **f**) compared to the original library (**e**). Each dot represents the average of tested codons for an amino acid. **h**, Flow cytometry plot of BW242 binding and HIT expression in T cells edited with a HIT β112 saturation mutagenesis pool. Boxes indicate sorted populations. **i**, Relative enrichment of mutations in HIT^+^ BW242^-^ cells versus HIT^+^ BW242^+^ cells. Each dot represents enrichment for a single codon. Lines display the average enrichment of all codons for an amino acid. **j,** BW242 binding for HIT^+^ T cells edited with individual HIT variants (*n* = 3 donors) compared with average enrichment scores (from **i**). Black line indicates BW242 MFI for TCR^-^ cells. MFI SEM displayed by bars (HIT mutants) or shaded grey area (TCR^-^). **k**, Diagram of a *TRAC* intron-targeted SEED HDRT encoding an epitope-edited HIT. **l,m**, T cells were edited with a HIT SEED and then immunomagnetically purified with BW242 (*n =* 3 donors) **l,** Flow cytometry plots of BW242 binding and HIT expression **m,** Percentage of BW242^+^ and BW242*^-^* HIT^+^ cells. **n**, Cytotoxic activity of T cells with a non-modified HIT (blue), epitope-edited HIT (red), or TCR knockout (grey) against Nalm6 lines (*n* = 1 donor, technical triplicate). Histograms show Nalm6 CD19 expression (flow cytometry). Unstained cells shown in grey. Bars display SEM. BP: branchpoint; SA: splice acceptor; SD: splice donor; LHA/RHA: homology arms; MFI: median fluorescence intensity; SEM: Standard Error of the Mean.

Previous studies have established that certain α/β TCR-targeted antibody epitopes can be disrupted by murinizing a portion of the β constant domain (Cβ) of a TCR^24,35^. To identify single amino acid (AA) mutations capable of disrupting TCR-targeted antibody binding, we systematically substituted individual residues across a 50 AA span of the HIT Cβ in the form of a 649-member pooled knock-in library (Fig. 2d,e). After introducing the receptor pool into the *TRAC* locus of T cells, we immunomagnetically depleted cells that were bound by a GMP-grade anti-TCR antibody (clone BW242/412 – hereafter referred to as BW242) and sorted for cells that retained high HIT expression (Fig. 2f). Library member abundance was then quantified using RNA-based next-generation sequencing (NGS) (Fig. 2g)^8^. While most library members were depleted, a subset of substitutions at Cβ residues G102, D112, or P116 were enriched, suggesting that these positions interact with BW242 and can be mutated to prevent HIT receptor depletion without compromising surface expression.

To catalogue conservative and non-conservative substitutions at these key positions, we generated three pooled libraries of HIT receptors with saturation mutagenesis performed at either Cβ 102, 112, or 116. Each library was individually introduced into T cells, and HIT^+^ cells were sorted based on BW242 binding (Fig. 2h and Extended Data Fig. 5a). RNA-based NGS was then used to assess the relative enrichment of mutants in each bin as compared to the original library. Residues at each position were ranked least-to-most conservative based on the ratio of enrichment between bins (Fig. 2i and Extended Data Fig. 5b). As expected, the native Cβ residue (G102, D112, P116) at each position was the most conservative. Almost all substitutions at 102 and 116 were preferentially enriched in the BW242^-^ bin, suggesting that the native residues at these positions are required for optimal BW242 binding (Extended Data Fig. 5b). 14 of 19 substitutions at Cβ 112 were also enriched in the BW242^-^ bin (Fig. 2i). However, substitutions to E, N, G, P, and W were preferentially enriched in the BW242^+^ bin. The most conservative substitution at Cβ 112 preserved the charge (D>E) whereas the least conservative ones switched the charge (D>K; D>R), suggesting that electrostatic interactions with D112 contribute to BW242 binding.

To confirm that this sequencing-based approach for epitope mapping predicts functional changes in antibody binding, we generated five HIT receptors with different residues at Cβ 112 and individually introduced them into T cells. BW242 binding was assessed by flow cytometry and compared to the enrichment ratios obtained from the Cβ 112 saturation mutagenesis screen (Fig. 2j and Extended Data Fig. 5c). Sequencing-based epitope mapping correctly ranked the panel of substitutions based on epitope conservation and was able to accurately detect variation in BW242 binding across the dynamic range. HIT and CD3 surface expression was consistent across receptor designs, confirming that variations in BW242 binding were not due to alterations in HIT expression or assembly with CD3 (Extended Data Fig. 5d). Of the five tested mutants, HIT 112K had the least BW242 binding and was selected for functional characterization (Fig. 2j).

We introduced a CD19-specific HIT^112K^ into a *TRAC*-targeted SEED and confirmed that HIT^+^ cells could be enriched by TCR depletion using BW242, demonstrating successful removal of all endogenous TCRs from the population while preserving an ultra-high purity (98%) of HIT^+^ cells (Fig. 2k–m). As the HIT receptor was developed to have the capacity to target tumor cells expressing low antigen densities^31^, we assessed receptor function through cytotoxicity assays with target cell lines expressing high or low levels of CD19 (Fig. 2n). HIT^112K^ performed equivalently to non-modified HIT (HIT^WT^) against high and low antigen density lines, confirming that epitope editing allows HIT^+^ cells to be enriched through SEED-Selection without compromising receptor function.

### Epitope editing allows for the identification and removal of *TRAC*-edited TCR-reprogrammed cells that exhibit mispairing

T cells can be reprogrammed to target a specified peptide-MHC through the expression of a transgenic TCR^30^. However, mispairing between endogenous and transgenic TCR chains can occur when a transgenic TCR is expressed in an otherwise unedited T cell^24,36^. Since all TCR chains compete for the same pool of CD3 molecules during assembly, mispairing can reduce the surface expression of the correctly paired transgenic TCR^24,36^. Additionally, T cells with mispaired TCRs have unknown specificities and can target healthy tissue leading to toxicities such as graft-versus-host disease (GVHD)^36^.

Mispairing is a non-random process influenced by properties of the transgenic TCR as well as each T cell’s endogenous TCR, and some combinations of TCR chains pair inefficiently^37^. In *TRAC*-edited cells engineered to express a transgenic TCR, mispairing occurs between the transgenic TCRα and endogenous TCRβ^24^. Therefore, we hypothesized that editing the BW242 epitope on a transgenic TCR β chain could allow cells that expressed a correctly paired transgenic TCR (which should evade BW242 binding) to be distinguished from cells that expressed a mispaired TCR (which should retain BW242 binding) (Fig. 3a).

**Figure 3:**
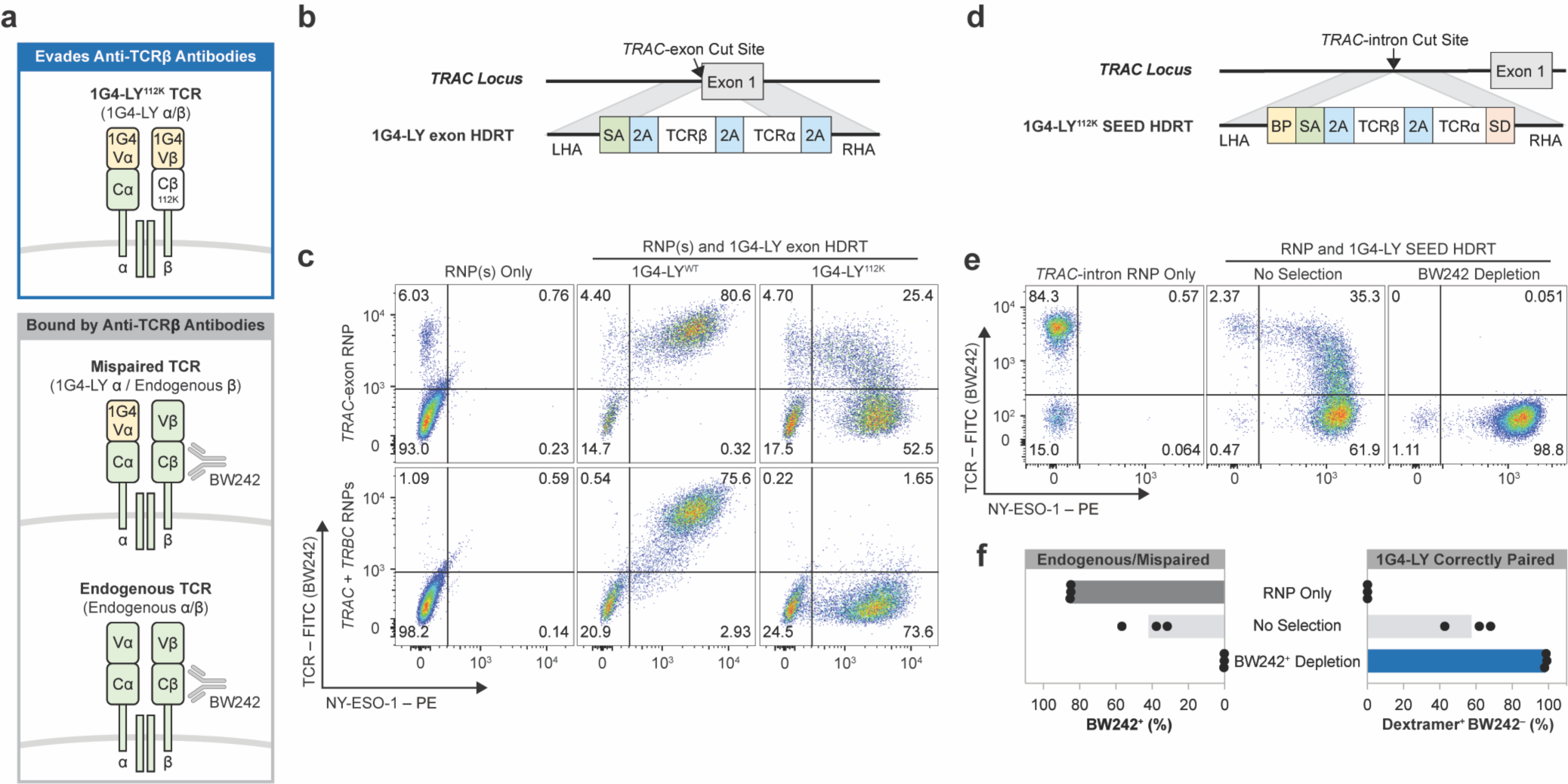
SEED-Selection depletes *TRAC*-edited TCR-swapped cells that express mispaired TCRs. **a,** Schematic of possible TCR pairs in non-edited and *TRAC*-edited T cells engineered to express the 1G4-LY^112K^ TCR (specific for NY-ESO-1). **b,** Diagram of a *TRAC* exon-targeted HDRT encoding 1G4-LY. **c,** Flow cytometry plots of BW242 binding and NY-ESO-1 dextramer binding in CD8^+^ cells. Editing was performed with RNP(s) targeting *TRAC* or *TRAC/TRBC* and HDRTs (**a**) encoding non-modified (1G4-LY^WT^) or epitope-edited (1G4-LY^112K^) versions of 1G4-LY (*n =* 2 donors). **d**, Diagram of a *TRAC* intron-targeted SEED HDRT encoding 1G4-LY. **e,f,** T cells were edited with a 1G4-LY SEED (**d**) and immunomagnetically purified with BW242 (*n* = 3 donors). **e,** Flow cytometry plots of BW242 binding and NY-ESO-1 dextramer binding for cells gated on CD8^+^. **f,** Percentage of cells with an endogenous or mispaired TCRs (BW242^+^) or correctly paired 1G4-LY (Dextramer^+^ BW242*^-^*). BP: branchpoint; SA: splice acceptor; SD: splice donor; LHA/RHA: homology arms.

To test this hypothesis, we created an epitope-edited variant of 1G4-LY: a clinically-validated, affinity-matured, transgenic TCR that targets NY-ESO-1 (SLLMWITQC) on HLA-A*02^38^. We designed *TRAC* exon-targeted HDRTs to simultaneously disrupt the endogenous TCRα and facilitate expression of a non-modified 1G4-LY (1G4-LY^WT^) or an epitope-edited variant (1G4-LY^112K^) (Fig. 3b). Each HDRT was introduced into CD8^+^ T cells edited with an RNP targeting *TRAC*. Multiplexed editing of *TRAC* and *TRBC* was also performed to create T cells where 1G4-LY was expressed and both endogenous TCR chains were disrupted, ensuring exclusive expression of correctly paired 1G4-LY.

NY-ESO-1 dextramer binding correlated with BW242 binding in cells expressing 1G4-LY^WT^, as expected (Fig. 3c). In contrast, *TRAC*-edited cells engineered to express 1G4-LY^112K^ exhibited a distinct population with correct TCR pairing (dextramer^+^ BW242^-^) and a separate population with apparent TCR mispairing (dextramer^+^ BW242^+^). In agreement, this population was eliminated by concurrent knockout of the endogenous TCRβ chain (Fig. 3c). Dextramer^+^ cells in all conditions expressed similar levels of CD3, confirming that variations in BW242 binding were not caused by differences in TCR assembly (Extended Data Fig. 6a). We also used epitope editing to identify and deplete mispaired *TRAC*-edited cells engineered to express a transgenic TCR (DMF5) targeting MART-1, supporting the generalizability of this approach (Extended Data Fig. 6b,c)^39^.

To test whether an epitope-edited TCR could be enriched through SEED-Selection, we introduced 1G4-LY^112K^ into a *TRAC*-targeted SEED and edited CD3^+^ T cells (Fig. 3d). Cells that expressed endogenous or mispaired TCRs were efficiently depleted (<1% BW242^+^) through immunomagnetic selection with BW242, leaving a highly pure (>98%) population of 1G4-LY^+^ cells with minimal detectable mispairing (Fig. 3e,f). This result suggests that SEED-Selection could be used as a simple method to deplete T cells with undesired specificities without the need to perform simultaneous editing at *TRAC* and *TRBC*.

### Multiplexed SEED-Selection enables the production of hypoimmune, co-receptor swapped, TCR-swapped cells with minimal mispairing

Recent studies have emphasized the role of CD4^+^ T cells as contributors to long-lasting immune responses to tumors^40^. However, the MHC class I–restricted TCRs commonly isolated for transgenic TCR therapies perform sub-optimally in the absence of the CD8α/β co-receptor^41,42,43,^. Although 1G4-LY has been demonstrated to undergo co-receptor independent activation^38^, we observed that dextramer binding was markedly reduced in edited CD4^+^ T cells as compared to edited CD8^+^ T cells (Extended Data Fig. 7a). Therefore, we sought to develop a strategy for isolating CD4^+^ T cells edited to express both CD8α/β and 1G4-LY.

Since CD4 should be superfluous in cells engineered to express an MHC class I–restricted TCR, we screened for non-disruptive gRNAs targeting the *CD4* locus and designed a SEED HDRT to deliver CD8α and CD8β (Fig. 4a and Extended Data Fig. 7b). Integration-mediated CD4 disruption (CD4^-^ CD8^+^) was achieved in >75% of HDRT-transduced CD4^+^ cells (Fig. 4c). Full disruption of CD4 requires both alleles to be non-functional. Correspondingly, a population that co-expressed intermediate levels of CD4 and CD8 was observed after editing, consistent with monoallelic SEED integration (Extended Data Fig. 7c). Cells with endogenous and intermediate CD4 expression were efficiently depleted through immunomagnetic selection, leaving a highly pure (>98%) population of biallelically edited (CD4^-^ CD8^+^) cells (Fig. 4c).

**Figure 4:**
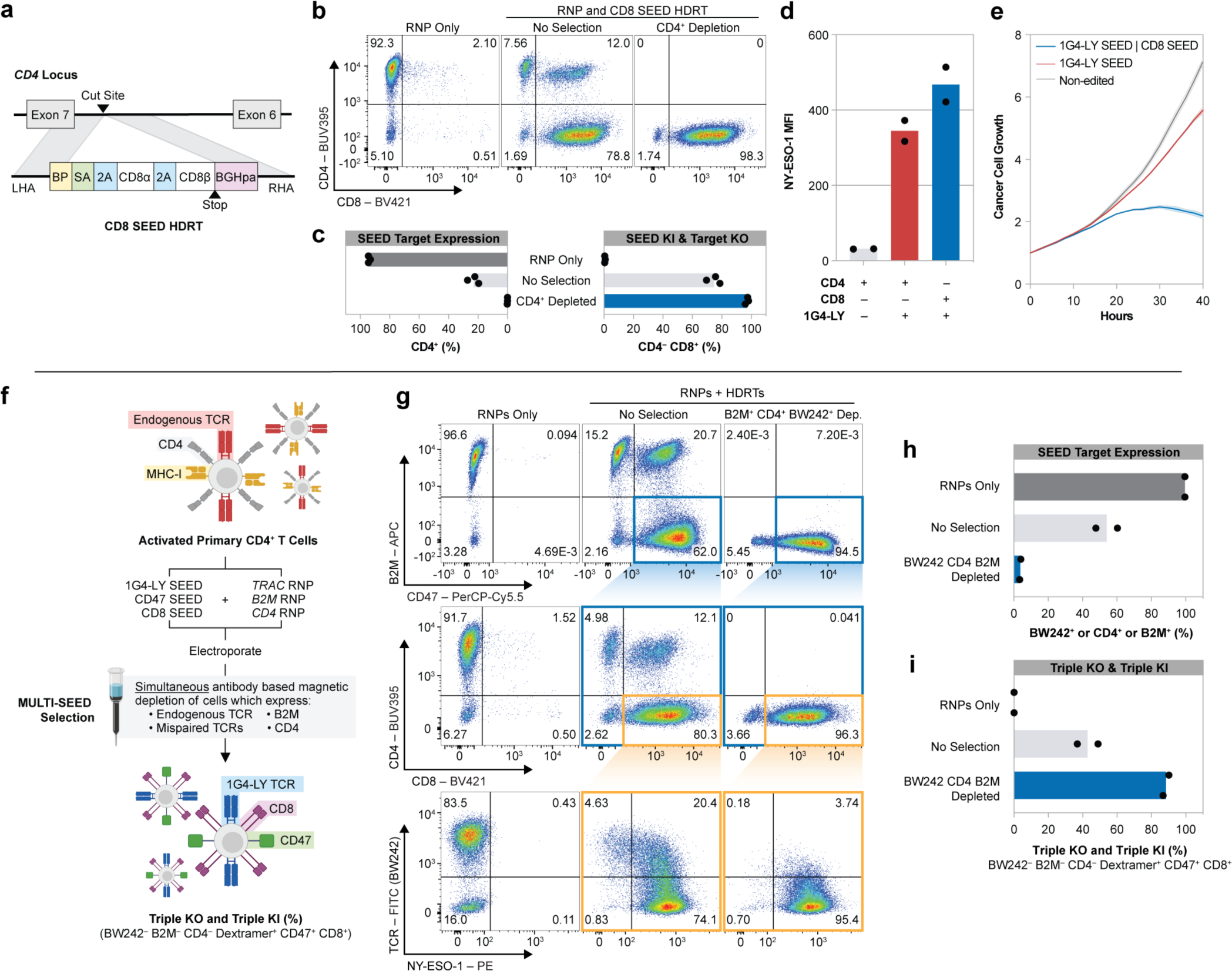
SEED-Selection enables the enrichment of TCR-swapped, co-receptor-swapped hypo-immune cells. **a,** Diagram of a *CD4* intron-targeted SEED HDRT encoding CD8α/β. **b,c**, CD4^+^ T cells were edited with a CD8 SEED (**a**) and immunomagnetically purified with anti-CD4 (*n* = 3 donors). **b,** Flow cytometry plots of CD4 and CD8 expression. **c,** Percentage of CD4^+^ and CD4^-^ CD8^+^ cells. **d,** Assessment of NY-ESO-1 dextramer binding by flow cytometry in CD4^+^ CD8^-^ BW242^-^ 1G4-LY^+^ cells (red) and CD4^-^ CD8^+^ BW242^-^ 1G4-LY^+^ cells (blue) (*n* = 2 donors, technical triplicate). **e**, Representative assessment of cytotoxic activity of CD4^+^ T cells against NY-ESO-1^+^ A375 target cells seeded at an initial 1:2 effector:target ratio. Non-edited T cells (dark gray), 1G4-LY SEED edited (red), 1G4-LY SEED and CD8 SEED edited (blue). Shaded areas indicate SEM (*n* = 2 donors). **f–i**, CD4^+^ T cells were edited with RNPs targeting *TRAC*, *B2M*, and *CD4* and transduced with SEEDs encoding 1G4-LY, CD8, and CD47, respectively. Edited cells were then immunomagnetically purified with BW242, anti-CD4, and anti-B2M *(n =* 2 donors). **f,** Diagram of the workflow for multiplexed editing and enrichment. **g,** Flow cytometry plots of SEED target expression (endogenous TCR, CD4, B2M), SEED payload expression (CD8, CD47), and NY-ESO-1 dextramer binding. Colored boxes indicate subpopulations. **h**, Percentage of cells expressing any SEED target or a mispaired TCR (BW242^+^ or CD4^+^ or B2M^+^). **i,** Percentage of triple knockout/triple knock-in cells with correct 1G4-LY pairing (BW242^-^ B2M^-^ CD4^-^ Dextramer^+^ CD47^+^ CD8^+^). BP: branchpoint; SA: splice acceptor; BGHpa: bovine growth hormone polyA; SD: splice donor; LHA/RHA: homology arms. SEM: Standard Error of the Mean

To assess whether the performance of 1G4-LY in CD4^+^ cells was improved through overexpression of CD8α/β, we edited T cells with a *TRAC*-targeted 1G4-LY^112K^ SEED alone or with a *CD4*-targeted CD8α/β SEED. Each cell population was immunomagnetically purified with antibodies targeting TCR (BW242) and CD4 to enrich for transgene integration and to deplete cells with mispaired and endogenous TCRs (Extended Data Fig. 7d). As expected, cells that expressed both 1G4-LY^112K^ and CD8α/β exhibited increased NY-ESO-1 dextramer binding in comparison to cells that expressed 1G4-LY^112K^ alone (Fig. 4d). Furthermore, in a longitudinal cytotoxicity assay, co-expression of CD8α/β improved the control of A375, a melanoma line that endogenously expresses NY-ESO-1 (Fig. 4e).

To validate the ability of SEED-Selection to isolate fully edited cells after complex editing, we sought to simultaneously select for cells with clinically desirable transgenes integrated at three genomic loci (Fig. 4f). CD4^+^ cells were edited with RNPs targeting *TRAC*, *B2M*, and *CD4* and transduced with SEEDs encoding IG4-LY^112K^, CD47, and CD8, respectively. After editing, cells expressing any combination of endogenous TCR, mispaired TCR, B2M, or CD4 were immunomagnetically removed in a single step. Depletion of all target markers was efficient, resulting in the isolation of highly pure (up to 90%) populations of fully edited BW242^-^ B2M^-^ CD4^-^ 1G4-LY^+^ CD47^+^ CD8^+^ cells (Fig. 4g–i and Extended Data Fig. 8). These findings demonstrate a one-step method for negative selection of cells with complex editing for reprogrammed specificity and function.

## Discussion

SEED-Selection has many characteristics that are desirable for clinical applications (Fig. 5). The immunomagnetic reagents used in SEED-Selection are highly amenable to automation and already used in manufacturing workflows. Additionally, SEED-Selection does not require the expression of exogenous proteins such as drug resistance cassettes, which can provoke an immune response^44^.

**Figure 5:**
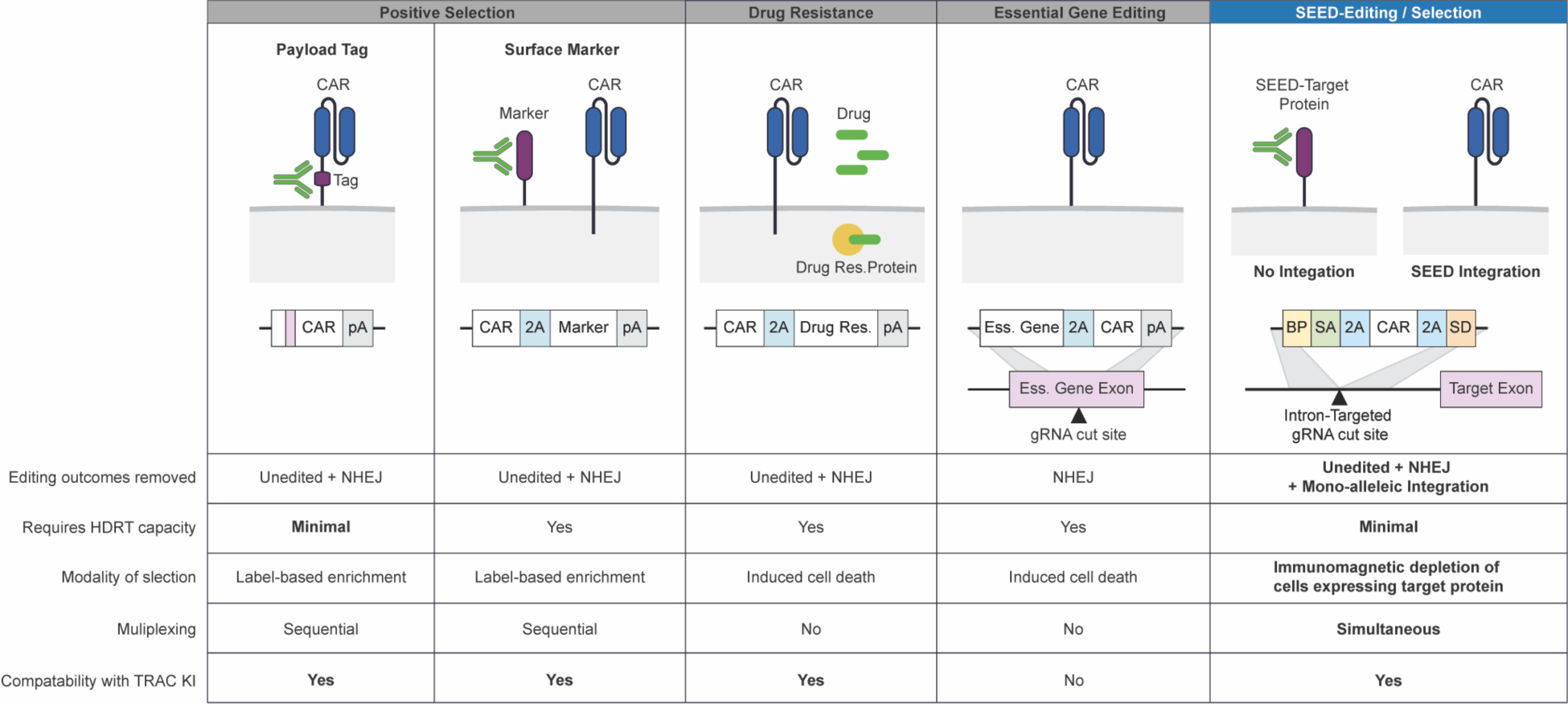
SEED-Selection compared to existing enrichment strategies. Chart detailing different strategies for isolating cells engineered to express a CAR or other transgene. Representative HDRTs are shown for each selection strategy. BP: branchpoint; SA: splice acceptor; SD: splice donor; pA: polyA signal; LHA/RHA: homology arms.

The reductive nature of SEED-Selection leaves isolated cells unlabeled and allows for multiple SEEDs to be simultaneously enriched. While we demonstrate that up to three SEEDs can be enriched in a single round of selection, immunomagnetic panels targeting 10 or more surface proteins are routinely used in laboratory and clinical settings to isolate rare (<1% of total cells) lymphocyte subsets from blood^45^. Therefore, we envision that more complex SEED-Selection strategies could be implemented as genome editing technologies advance.

Although we initially tested SEEDs with AAV delivery, non-viral DNA delivery vehicles such as linear ssDNA, dsDNA, or nanoplasmid could be used^11,14,46^. Additionally, SEED-Selection could be used to enrich for targeted transgene integration in other clinically relevant cell types, such as hematopoietic stem cells, induced pluripotent stem cells, or NK cells.

Chromosomal rearrangements such as translocation and chromosomal loss have been observed in gene-edited primary cells^47–49^. While we show that SEED-Selection can be used to deplete cells with undesirable editing outcomes (repair through NHEJ and monoallelic integration), further studies are necessary to determine whether this process also depletes cells with other unintended editing outcomes. Additionally, SEED-Selection could be used with methods that allow edits at different loci to be performed sequentially, which has been shown to reduce the occurrence of translocations^6^.

In summary, SEED HDRTs and SEED-Selection provide a simple, one-step process for isolating highly pure populations of cells with multiple transgene integrations. We expect that this approach will streamline manufacturing for current products and enable the development of more advanced cellular therapies.

## Methods

### T cell isolation & culture

Primary adult blood cells from anonymous healthy human donors were purchased as leukapheresis packs (Stemcell) and cryopreserved. Specific lymphocytes were isolated from thawed aliquots using EasySep isolation kits for CD3^+^, CD4^+^, or CD8^+^ T cells (Stemcell). Isolated T cells were cultured at an initial density of 10^6^ cells/mL in X-Vivo 15 medium (Lonza) supplemented with human serum (5%, Gemini), Penicillin-Streptomycin (1%, Gibco), IL-7 (5 ng/mL, Miltenyi), and IL-15 (5 ng/mL, Miltenyi). After isolation, cells were stimulated for two days with anti-human CD3/CD28 magnetic Dynabeads (Thermo) using a 1:1 bead-to-cell ratio.

### NK cell isolation & culture

PBMCs were obtained through density gradient centrifugation of Trima residuals from apheresis collection (Vitalant). NK cells were isolated using EasySep Human NK Cell Enrichment Kits (Stemcell). Isolated NK cells were cultured at an initial density of 10^6^ cells/mL in NK MACS medium (Miltenyi) supplemented with human platelet lysate (5%, Elite Cell), Penicillin-Streptomycin (0.5%) and IL-2 (1000U/mL, Peprotech), as previously described^50^. Cells were stimulated for seven days with anti-human CD2/NKp46 beads (Miltenyi) using a 1:2 bead-to-cell ratio. After bead removal, cells were subsequently cultured and replated twice per week at 10^6^ cells/mL.

### HDRT Design

Sequences for individually tested HDRTs and site saturation mutagenesis pools are provided in Supplementary Table 1. Splicing elements at the 5’ end of SEEDs (which include the polypyrimidine tract, branchpoint sequence (BP), and splice acceptor (SA)) were derived from the chimeric intron included in the pCI mammalian expression vector (Promega). We used a P2A sequence to prematurely truncate the SEED-target and facilitate expression of a transgene payload. We selected integration sites between the signal peptide and transmembrane domain of the SEED-target, so that SEED-target surface expression would be disrupted upon HDRT integration. In most SEED designs, the 3’ end of the SEED included an additional P2A followed by the splice donor (SD) sequence of the preceding exon. This allows the transgene to be expressed with the endogenous polyA sequence of the SEED-target and conserves HDRT cargo capacity. In SEED HDRTs encoding transgenic TCRs or HITs, the 3’ P2A was excluded to allow the HIT/TCRα chain to be completed using the endogenous *TRAC* sequence. Alternatively, our *CD4*-targeted SEED did not include a final P2A or splice donor sequence and instead relied on a bovine growth hormone polyA (BGHpa) signal. Where necessary, additional nucleotides were added downstream of the SA and/or upstream of the SD to maintain the reading frame of the spliced transcript.

### AAV production

AAV plasmids were packaged into AAV6 by transfection of HEK293T cells and purified using iodixanol gradient ultracentrifugation. Titers were determined by quantitative PCR on DNase I (NEB) treated and proteinase K (Qiagen) digested AAV samples. HDRTs targeting *TRAC* exon 1 were quantified using primers targeting the left homology arm of the HDRT, while all other AAVs were quantified using primers targeting the inverted terminal repeat (ITR) sequences (Supplementary Table 2). Quantitative PCR was performed with SsoFast EvaGreen Supermix (Bio-Rad) on a StepOnePlus Real-Time PCR System (Applied Biosystems).

### RNP formulation

gRNA sequences are provided in Supplementary Table 3. For most experiments, RNP was generated by incubating single guide RNAs (Synthego) with Cas9 protein (40 µM, Berkeley QB3 MacroLab) at a 2:1 (sgRNA:Cas9) molar ratio for 15 minutes at 37 °C. For intron gRNA screening experiments, RNPs were produced by complexing a two-component gRNA (Edit-R, Dharmacon Horizon) to Cas9 protein with the addition of a PGA (Sigma) electroporation enhancer, as previously described^11^. When multiple loci were targeted, RNPs were individually complexed and then mixed shortly before electroporation.

### T cell editing

For each electroporation, 2×10^6^ cells were resuspended in P3 buffer (Lonza), mixed with RNP(s), and added to a 96-well nucleofection plate (Lonza). RNP amounts per electroporation varied based on the number of loci targeted: (1 locus) 60 pmol of RNP; (2 loci) 60 pmol of each RNP, 120 pmol total; (3 loci) 53 pmol of each RNP, 159 pmol total. P3 buffer volume was adjusted so that the total volume of each reaction was 23 μL.

Cells were electroporated using a Lonza 4D-Nucleofector 96-well unit (Code: EH-115). Pre-warmed X-Vivo 15 medium (<u>without</u> human serum) was then added to achieve a density of 2×10^6^ live cells/mL, assuming a one-third loss of viability after electroporation. AAV6 encoding HDRT(s) was added to cultures shortly after editing. For most experiments, a multiplicity of infection of 2×10^5^ was used. After an overnight incubation, edited cells were resuspended in fresh complete medium. Edited cells were subsequently expanded, keeping a density of 10^6^ cells/mL.

### Flow cytometry/sorting

Flow cytometry was performed on a BD FACSymphony Fortessa X-50 or an Attune NxT. Cell sorting was performed on a BD FACSAria. Cells were resuspended in FACS buffer (phosphate-buffered saline (PBS), 2% FBS, and 1 mM EDTA) and stained with antibodies/dextramer (Supplementary Table 4). Zombie Violet (BioLegend) or Ghost Dye Red (Tonbo) were used in experiments where viability was assessed via flow cytometry. In experiments where HIT or CAR expression was assessed, cells were initially stained with anti-mouse F(ab’)_2_ and then blocked with mouse serum (MilliporeSigma) before antibody staining was performed. In experiments where anti-B2M and MHC-I dextramers were both used, cells were stained with antibodies first, washed, and then stained with dextramer. Flow cytometry analysis was performed in FlowJo (BD). Representative gating strategies are provided in the Supplementary Information.

### Intronic gRNA screening

Activated human T cells were edited with individual gRNAs and cultured for three days. Surface marker expression was then assessed via flow cytometry, and genomic DNA was isolated using QuickExtract (Epicenter). PCR amplification of cut site regions was performed with KAPA HiFi polymerase (Kapa Biosystems) according to the manufacturer-provided protocol. Amplicons were purified using SPRI beads (Beckman) and Sanger sequenced (Quintara Biosciences). The resulting sequencing files were aligned for detection of insertions and deletions using the ICE Analysis webtool (Synthego, https://ice.synthego.com/#/).

### SEED engineering/selection

For experiments with SEED HDRTs, M3814 (1μM, ChemieTek) was added to the recovery media after editing, unless otherwise specified. All purifications were performed 7–10 days after editing. Prior to selection, cell density and viability were assessed using a Countess II automated cell counter (Thermo Fisher). Cells were then centrifuged, resuspended in MACS buffer (80 μL per 10^7^ cells | PBS, 0.5M EDTA, 2% BSA), and incubated with a biotin-conjugated antibody (Supplementary Table 4) targeting the marker of interest (20 μL per 10^7^ cells) for 10 minutes at 4°C. In experiments where multiple markers were simultaneously depleted, cells were incubated with a master mix of antibodies (20 μL of each antibody per 10^7^ cells) and MACS buffer volume was adjusted to maintain a consistent incubation volume of (100 μL per 10^7^ cells). Cells were then washed, resuspended in MACS buffer (80 μL per 10^7^ cells), and incubated with anti-biotin microbeads (20 μL per 10^7^ cells, Miltenyi) for 15 minutes at 4°C. Labeled cells were loaded onto Miltenyi MACS columns and processed according to the manufacturer-provided protocol. Cell density in the flow-through from the column was assessed, and isolated cells were centrifuged and resuspended in complete T cell medium for culture. SEED-target and transgene payload expression was evaluated within 24 hours using flow cytometry.

### *B2M* integration site genomic DNA PCR

T cells were edited with *B2M* intron-targeted RNP (i4) and SEED HDRT encoding CD47. Edited cells were then expanded for seven days and immunomagnetically purified with anti-B2M antibody. Genomic DNA was isolated from non-edited cells, non-purified edited cells, and purified edited cells using a NucleoSpin Tissue Kit (Macherey-Nagel). The HDRT integration site was PCR-amplified from genomic DNA using Q5 High-Fidelity Polymerase (NEB), with an expected amplicon size of ∼1kb for non-edited *B2M* alleles and ∼2kb for alleles with HDRT integration. Primers were designed to target sequences upstream and downstream of the HDRT homology arms to avoid amplification of non-integrated repair template. Amplicon size was assessed using gel electrophoresis (0.8% agarose, 125V for 50 min) with SYBR Safe DNA stain (Thermo) using the 1kb Plus DNA ladder (NEB). Gel imaging was performed on a FluorChem M System (Cell Biosciences). Unprocessed gel images are provided in the Supplementary Information.

### HIT scanning mutagenesis library design and synthesis

A pooled library of oligos encoding Cβ residues 101–150 was designed to test substitutions at individual positions. To strike a compromise between library size and library diversity, we systemically chose substitutions using two processes: (1) At each residue, we substituted alanine along with four other substitutions that were predicted to be less disruptive based on scores from the BLOSUM80 matrix. Five substitutions from the BLOSUM80 matrix were introduced into residues that were originally alanine (Supplementary Table 5); (2) We aligned 38 homologous protein sequences (Supplementary Table 6) from mammalian species with the sequence for human TRBC. Homologous substitutions at a given position that were not already in the library were added.

Where possible, for each substitution, we synthesized two oligos with different codons. As a control, we also tested a single synonymous mutation at each position. Other controls included single-residue deletions (tested at all residues) and stop codons (tested at five individual residues within the mutagenesis region), resulting in a final library size of 649 oligos (Supplementary Table 7). The oligo pool was synthesized (Twist), PCR-amplified, and introduced into a plasmid backbone containing a *TRAC* exon-targeted HIT HDRT through Golden Gate Assembly.

### HIT site saturation mutagenesis library design and synthesis

gBlocks (IDT) encoding Cβ residues 101–150 were separately synthesized with degenerate nucleotides (N) specified at bases encoding a target residue (G102, D112, or P116). Each gBlock was PCR-amplified and individually introduced into a plasmid backbone containing a *TRAC* exon-targeted HIT HDRT through Golden Gate Assembly.

### HIT scanning mutagenesis screen

T cells were edited with *TRAC*-targeted RNP (i1) and AAV6 encoding the pooled HIT library. Edited cells were expanded, immunomagnetically purified with BW242, and sorted based on BW242 binding and HIT expression (using anti-mouse F(ab’)_2_). Bulk RNA was isolated from sorted cells and cells with the original library (Direct-zol RNA Microprep Kit, Zymo) and used as a template for cDNA synthesis (High-Capacity cDNA Reverse Transcription Kit, Applied Biosystems).

### HIT site saturation mutagenesis screen

T cells were edited with *TRAC*-targeted RNP (e1) and transduced with AAV6 encoding a single pooled HIT library (with site saturation mutagenesis performed at either Cβ 102, 112, or 116). Edited cells were expanded and sorted into two bins based on BW242 binding and HIT expression (using anti-mouse F(ab’)_2_). Bulk RNA was isolated from sorted cells and cells with the original library (Direct-zol RNA Microprep Kit, Zymo) and used as a template for cDNA synthesis (High-Capacity cDNA Reverse Transcription Kit, Applied Biosystems).

### Library amplification and analysis

cDNA from sorted and unsorted samples from all screens was processed using a shared workflow. PCR amplification of the library region was performed with Q5 High Fidelity Polymerase (NEB) using primers containing Illumina partial adapters. The resulting amplicons were purified using SPRI beads and submitted for 2 x 250bp paired-end next-generation sequencing (Amplicon-EZ, Genewiz/Azenta).

FASTQ files from NGS were processed using a workflow in Python. Briefly, reads were scanned for conserved sequences upstream and downstream of the library. Reads that contained these sequences (with no permitted mismatches) were trimmed so that only the library region remained, while reads that lacked these sequences were discarded. Trimmed reads were mapped to library members, with no permitted mismatches. Library member abundance within a given sample was calculated as: (# of reads mapped to library member) / (# of total mapped reads). Reads mapped to designated library controls (stop codons and deletions) were excluded from totals for analysis.

### HIT arrayed library screen

T cells were edited with *TRAC*-targeted RNP (e1) and separately transduced with HDRTs encoding a single HIT variant. After five days of expansion, HIT expression, CD3 expression, and BW242 binding were assessed via flow cytometry. Edited non-transduced cells were included as a control.

### Target cell culture

CD19^Hi^ and CD19^Lo^ Firefly luciferase^+^ Nalm6 cell lines were cultured in RPMI (Gibco) supplemented with FBS (10%), sodium pyruvate (1%, Gibco), HEPES buffer (1%, Corning), penicillin–streptomycin (1%), non-essential amino acids (1%, Gibco) and 2-mercaptoethanol (0.1%, Gibco). CD19 expression in Nalm6 lines was validated via flow cytometry. RFP^+^ A375 cells were cultured in Dulbecco’s modified Eagle medium (Gibco) supplemented with FBS (10%), sodium pyruvate (1%), HEPES buffer (1%) and penicillin– streptomycin (1%).

### CD47 SEED co-incubation with NK cells

A TCR^-^ population of T cells with (B2M^+^ CD47^-^; B2M^+^ CD47^+^; B2M^-^ CD47^+^) subsets was generated by performing editing with a *TRAC* exon-targeted RNP (e1), a *B2M* intron-targeted RNP (i4), and a CD47 HDRT without M3814. In parallel, TCR^-^ B2M^-^ CD47^-^ cells were generated by performing editing with a *TRAC* exon-targeted RNP (e1) and a *B2M* exon-targeted RNP (e1). Cells generated with both engineering strategies were immunomagnetically purified with BW242 to remove cells that retained TCR expression, and editing outcomes were assessed via flow cytometry. The two populations were then mixed to achieve an approximate 1:1 ratio of B2M^-^ CD47^-^ cells: B2M^-^ CD47^+^ cells. To block CD47, T cells were incubated with anti-CD47 antibody (Clone: B6H12, BD) for 30 minutes at 37°C and then washed with T cell medium. T cells were co-cultured overnight with allogeneic activated human NK cells, and then the co-culture composition was quantified via flow cytometry.

### CD19 cytotoxicity assay

T cells were co-cultured with 3×10^4^ Nalm6 cells in 96-well flat bottom plates. T cells were serially diluted (two-fold) from an initial 1:1 E:T ratio to a minimum 1:64 E:T ratio and plated in triplicate. Non-treated Nalm6 cells were included as a maximum signal control, and Nalm6 cells incubated with Tween-20 (0.2%) were included as a minimum signal control. After a 24-hour incubation, D-luciferin (0.75 mg/ml, GoldBio) was added to the plates, and luminescence was quantified using a GloMax Explorer microplate reader (Promega). Percentage cytotoxicity was determined as: 100% × (1 – (sample – minimum) / (maximum – minimum)).

### NY-ESO-1 cytotoxicity assay

T cells were co-cultured with 10^4^ pre-plated RFP^+^ A375 cells in 96-well flat bottom plates. T cells were serially diluted (two-fold) from an initial 2:1 E:T ratio to a minimum 1:16 E:T ratio and plated in triplicate. The RFP^+^ count per well was quantified every two hours over a 40-hour span using IncuCyte S3 live-cell imaging (Sartorius). A375 cell growth was calculated as the number of RFP^+^ objects at a given time point, normalized to the number of RFP^+^ objects at the start of the assay.

## Code Availability

Library analysis code will be provided by the corresponding author upon request.

## Data Availability

Sequences for all primers, guides, and HDRTs are provided in the supplementary dataset. Additional raw data will be provided by the corresponding author upon request.

## Acknowledgements

Schematics were created using elements from BioRender.com. We thank the PFCC (RRID:*SCR_018206*) for assistance generating flow cytometry data. Research reported here was supported in part by the DRC Center Grant NIH P30 DK063720. C.R.C. was supported by UCSF Medical Scientist Training Program (T32GM141323).

## Author Information

C.R.C., A.M., B.R.S, and J.E. conceived of and/or supervised the study. V.S.V. and B.R.S. screened gRNAs and designed HDRTs. C.R.C. designed and tested HDRTs, and designed and optimized immunomagnetic selection protocols. C.R.C, W.N., and C.L. designed and performed assays characterizing genomic editing outcomes. V.A. and A.R. contributed to functional assays relating to CD47. C.R.C. designed and performed the HIT epitope editing screen and characterized HIT mutants, with input from D.B.G. on pooled knock-in strategies. C.R.C designed and performed experiments characterizing and depleted cells with TCR mispairing. J.J.M., C.L, and C.H.W. contributed to virus production and quality control. C.R.C. and J.E. wrote the manuscript with input from all co-authors.

---

**Extended Data Figure 1:**
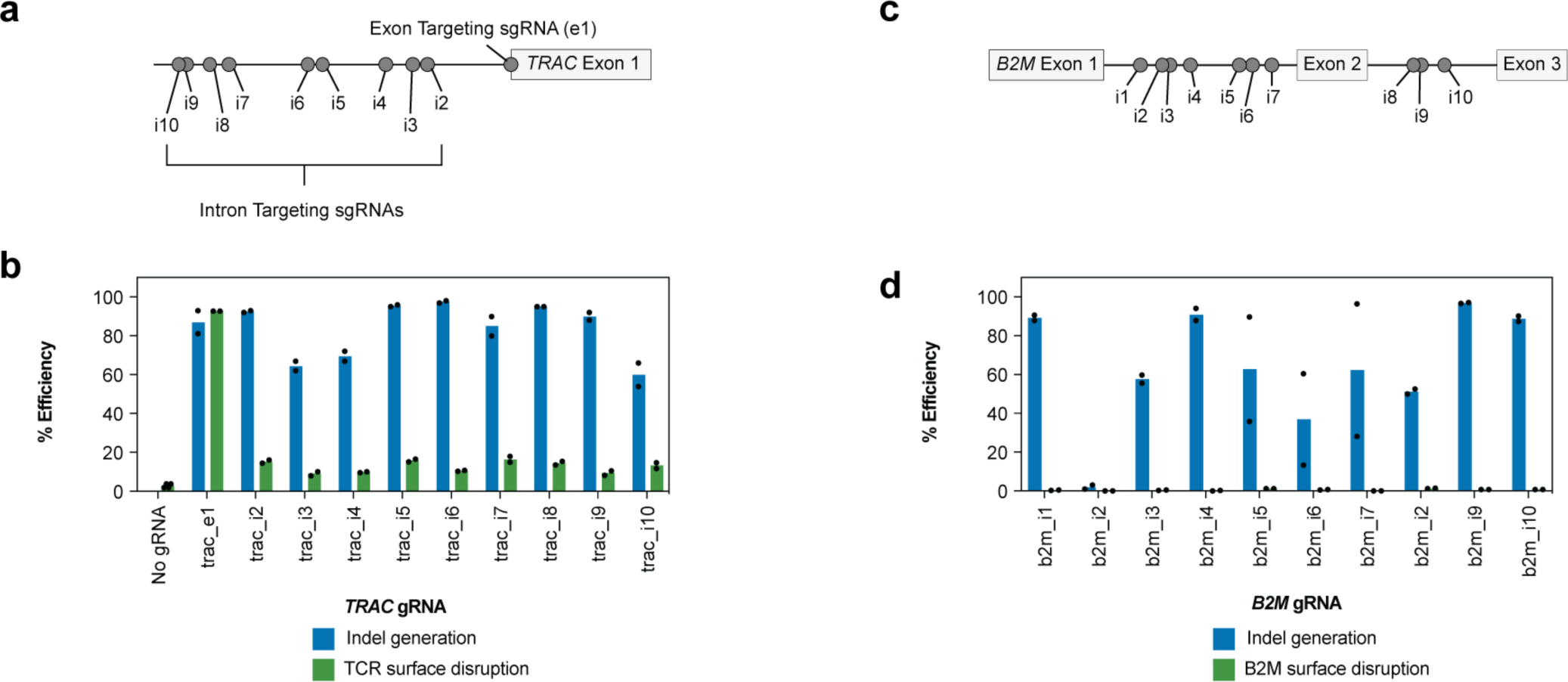
gRNA screens for *TRAC* and *B2M* introns. **a**, Panel of gRNAs targeting *TRAC* exon (e1) and *TRAC* intron (i2–i10); not depicted to scale. **b**, Assessment of TCR disruption by flow cytometry and indel generation by bulk genomic DNA sequencing in T cells after editing with gRNAs depicted in **b** (*n* = 2 donors). **c,** Panel of gRNAs targeting B2M introns; not depicted to scale. **d,** Assessment of B2M disruption by flow cytometry and indel generation by bulk genomic DNA sequencing in T cells after editing with gRNAs depicted in **c** (*n* = 2 donors).

**Extended Data Figure 2:**
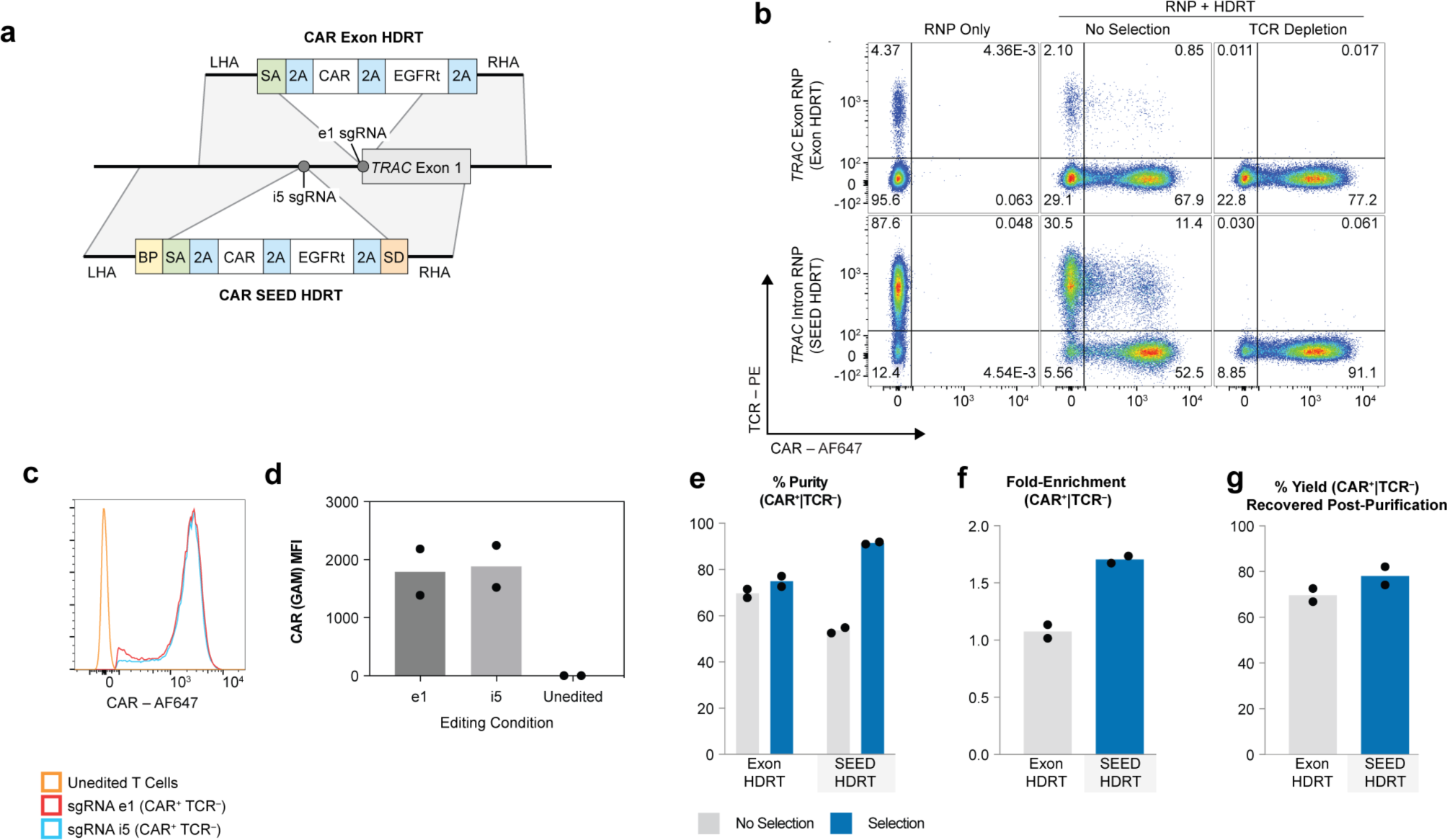
Intron-targeted gRNAs enable transgene integration through negative selection. **a**, Comparison of *TRAC*-intron and exon-targeted HDRTs encoding a CAR and EGFRt. **b–g**, T cells were edited with *TRAC* intron or *TRAC* exon-targeted RNPs and HDRTs (**a**). Edited cells from each condition were then immunomagnetically purified with anti-TCR (*n* = 2 donors). **b,** Flow cytometry plots of TCR and CAR (anti-mouse F(ab’)_2_) expression **c,** Histograms of CAR expression in TCR^-^ CAR^+^ cells. **d,** Median fluorescence intensity (MFI) of CAR expression in TCR^-^ CAR^+^ cells. **e**, Percentage of TCR^-^ CAR^+^ cells in purified (blue) and non-purified samples (grey). **f**, Relative enrichment of fully edited cells after purification: (% of purified sample) / (% of non-purified sample). **g**, Estimated percentage of fully edited cells recovered after TCR depletion, based on flow cytometry and cell counts: (# of fully edited cells after purification) / (# of fully edited cells set aside for purification).

**Extended Data Figure 3:**
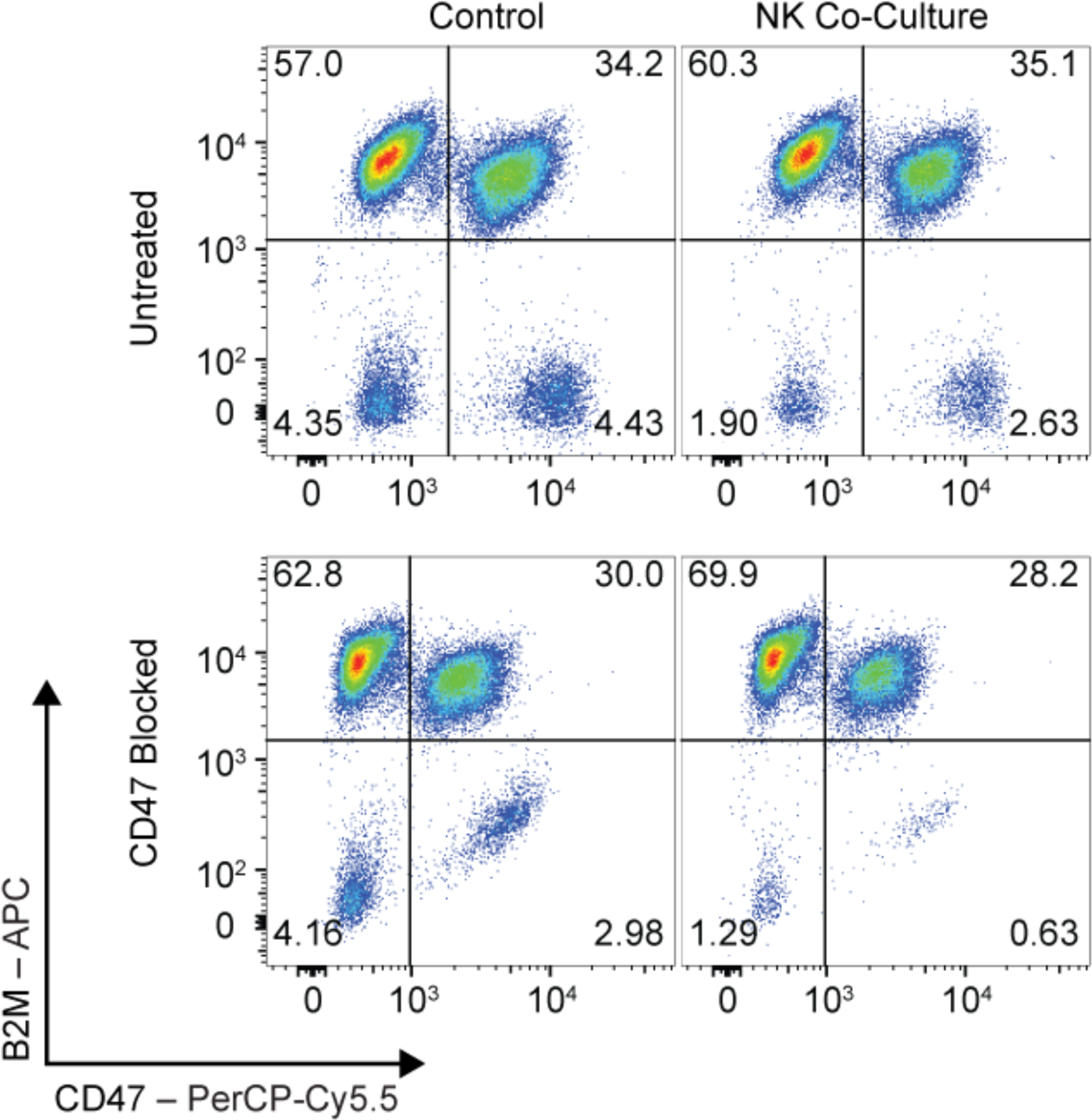
CD47 SEED integration reduces NK activity. Flow cytometry plots of B2M and CD47 expression in live CD56^-^ gated cells from edited T cells mixtures. Relative susceptibility to NK cells activity was determined by comparing the CD47^+^:CD47^-^ ratio in samples which were co-cultured with NK cells overnight to the CD47^+^:CD47^-^ ratio in a control sample. To block CD47 activity, cells were pre-treated with anti-CD47 (*n* = 2 donors).

**Extended Data Figure 4:**
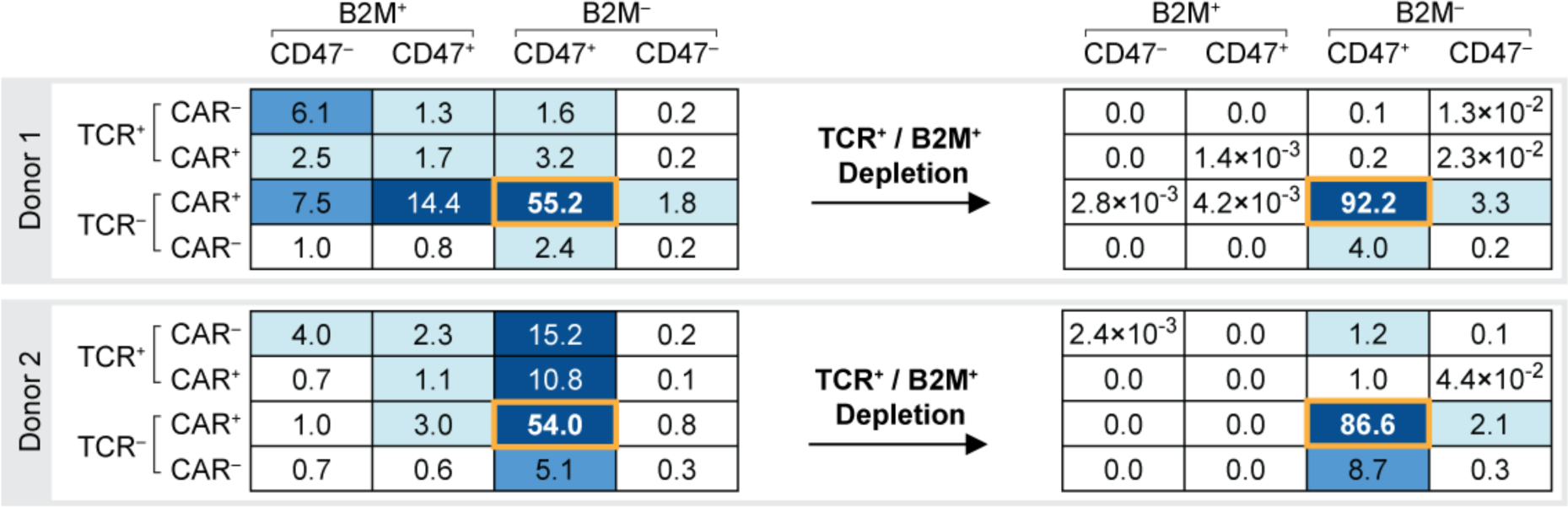
Outcomes for double SEED engineering and SEED-selection. Outcomes after multiplexed editing with CAR and CD47 SEED HDRTs and immunomagnetic purification with anti-TCR and anti-B2M. Numbers indicate percentage of total population of cells. Percentage of fully edited (TCR^-^ CAR^+^ B2M^-^ CD47^+^) in each condition is enclosed by an orange box.

**Extended Data Figure 5:**
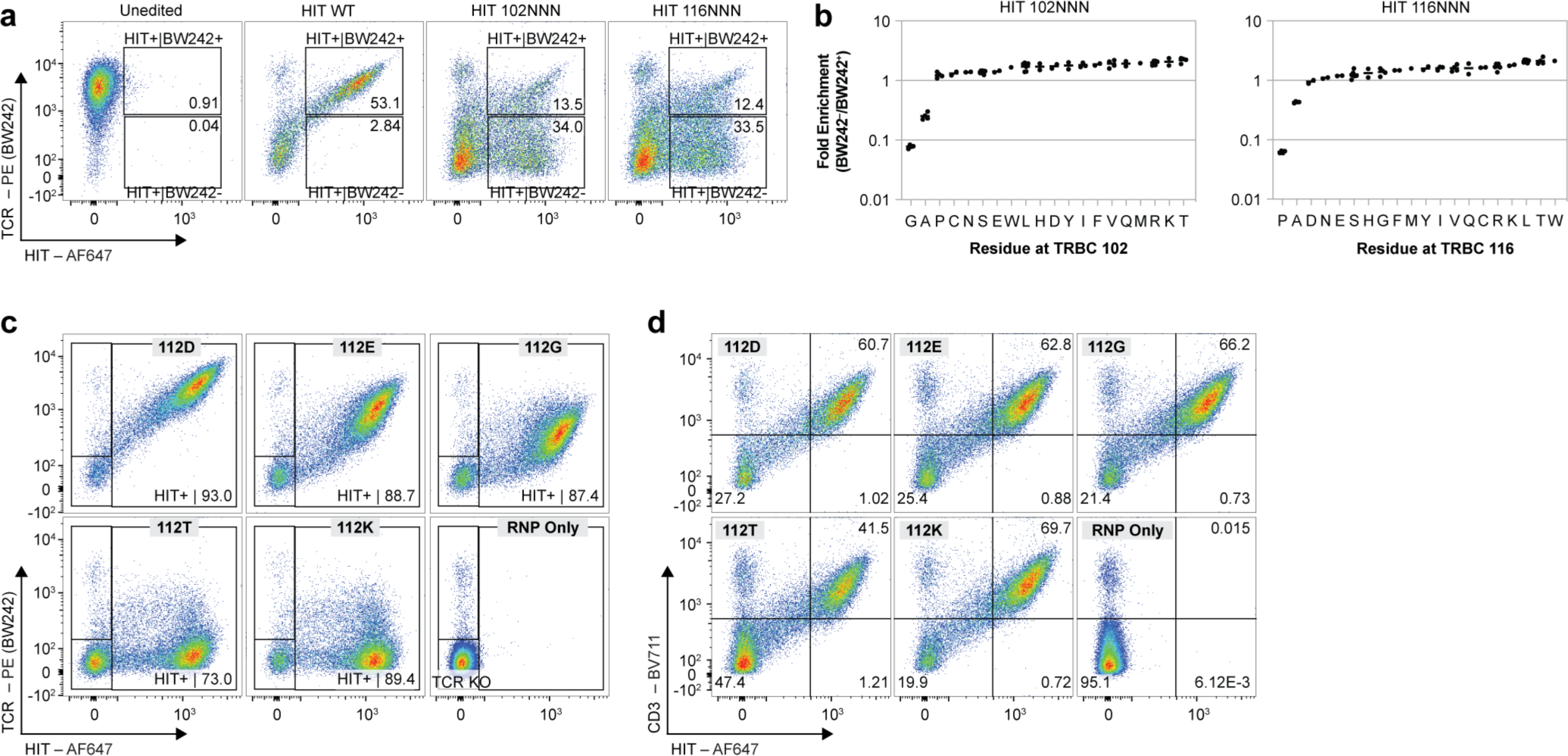
Characterization of individual and pooled libraries of epitope-edited HITs. **a,** Flow cytometry plots of BW242 binding and HIT expression (mouse F(ab’)_2_) in T cells edited with either a HIT β102 or β116 saturation mutagenesis pool. Boxes indicate sorted populations. **b**, Relative enrichment of mutations in HIT^+^ BW242^-^ cells versus HIT^+^ BW242^+^ in cells from **a**. Each dot represents enrichment for a single codon. Bars represent the average enrichment of all codons for an amino acid. **c,** Representative flow cytometry plots of BW242 binding and HIT expression in T cells individually edited with *TRAC*-exon targeted HDRTs encoding HIT receptor variants. Median fluorescence intensity (MFI) values for BW242 binding (for Fig. 2j) were determined based on the HIT^+^ gated population (for transduced samples) or the TCR KO gated population (*n =* 3 donors). **d,** Representative flow cytometry plots of CD3 and HIT expression in T cells individually edited with *TRAC*-exon targeted HDRTs encoding HIT receptor variants (*n* = 3 donors).

**Extended Data Figure 6:**
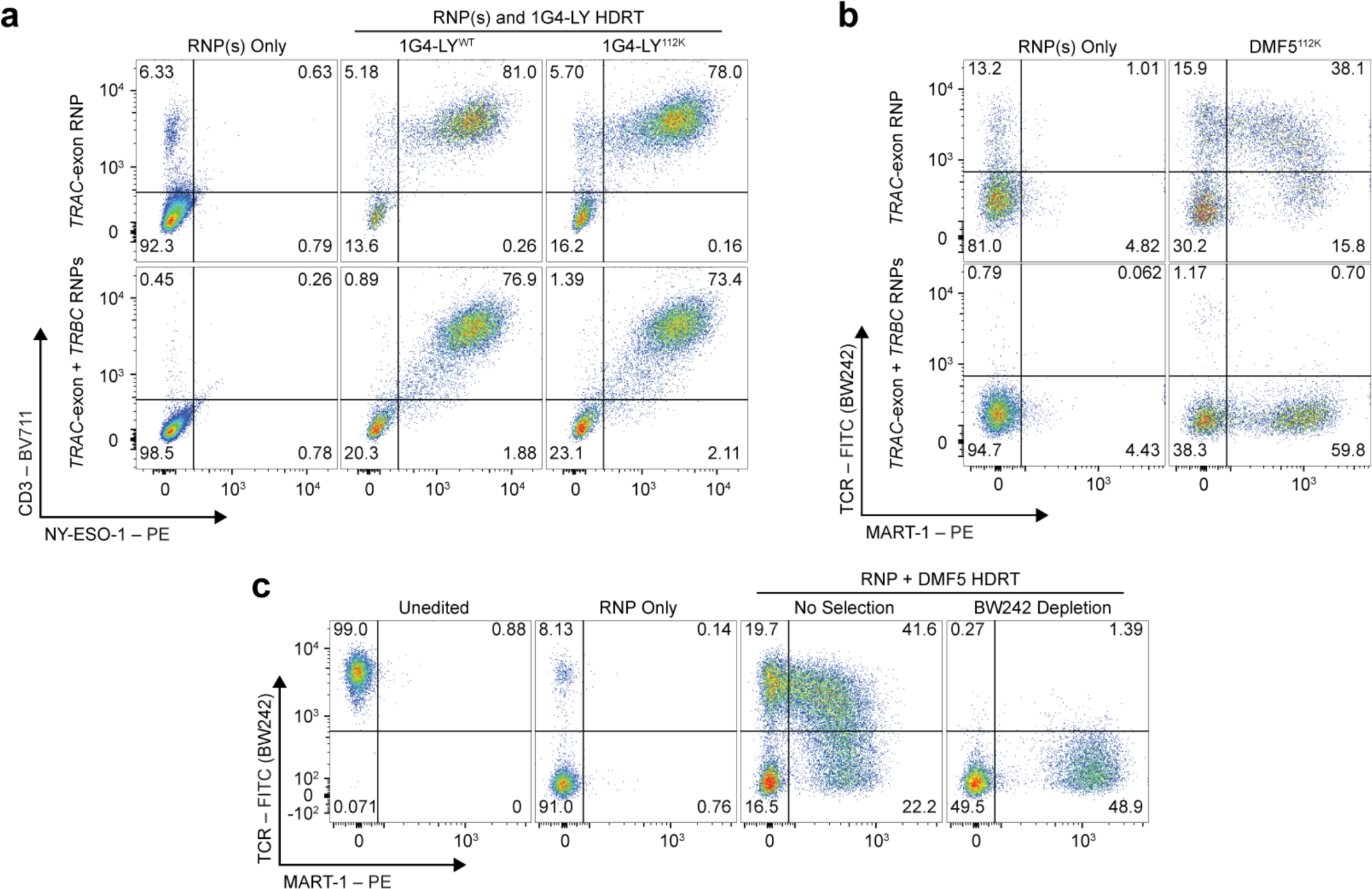
Validation of epitope editing in transgenic TCRs. **a**, Flow cytometry plots quantifying CD3 expression and NY-ESO-1 dextramer binding in CD8^+^ cells. Editing was performed with RNPs targeting *TRAC* or *TRAC/TRBC* and HDRTs encoding non-modified (1G4-LY^WT^) or epitope-edited (1G4-LY^112K^) versions of 1G4-LY (*n* = 2 donors). **b,** Flow cytometry plots quantifying BW242 binding and MART-1 dextramer binding in CD8*^+^* cells. Editing was performed with RNPs targeting *TRAC* or *TRAC/TRBC* and an HDRT encoding an epitope-edited version of a MART-1 TCR (DMF5) (*n* = 1 donor). **c,** Flow cytometry plots of BW242 binding and MART-1 dextramer binding in CD8^+^–gated T cells edited with a MART-1 TCR HDRT after immunomagnetic BW242 depletion (*n* = 1 donor).

**Extended Data Figure 7:**
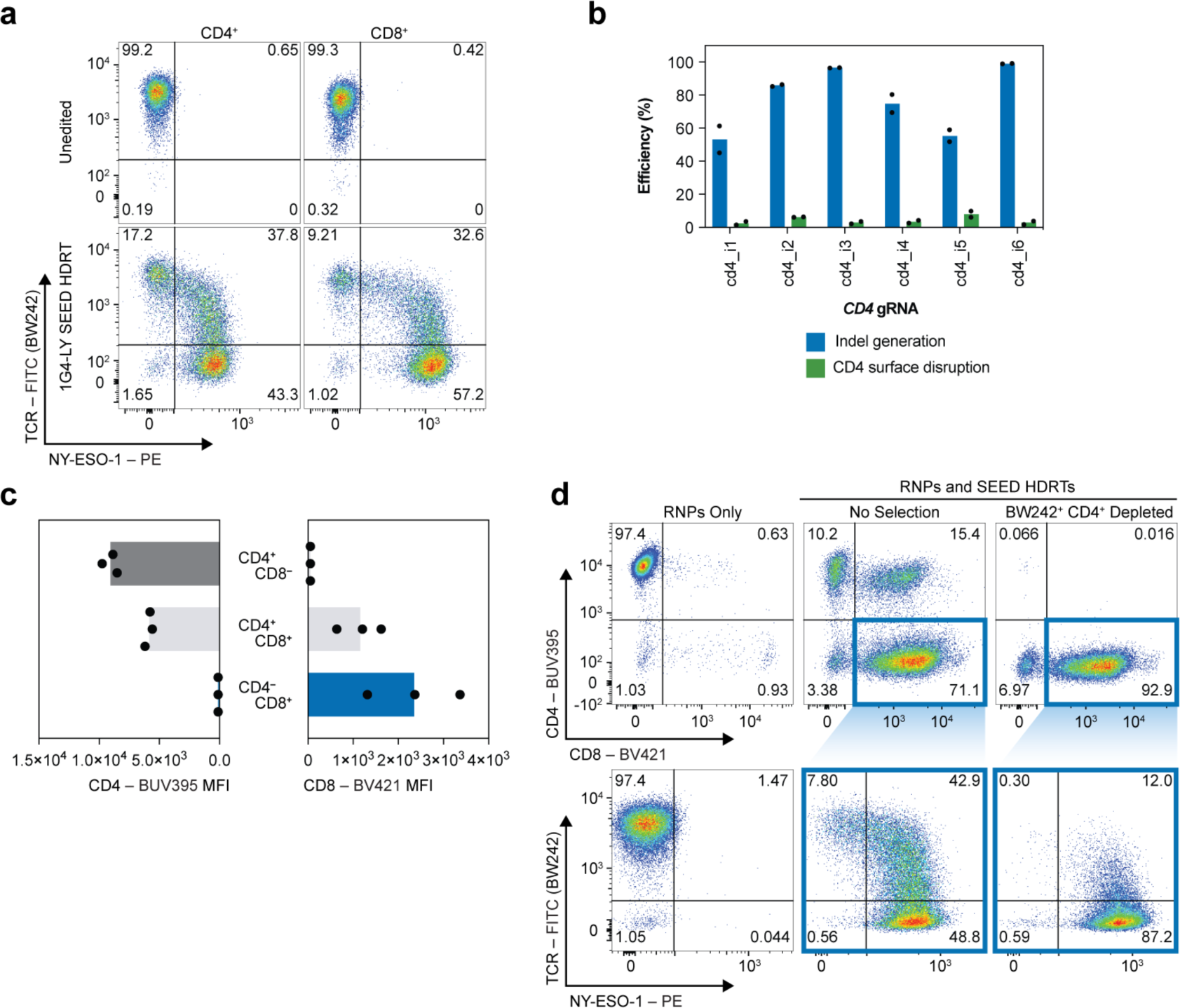
SEED-engineering facilitates co-receptor swapping. **a,** BW242 binding and NY-ESO-1 dextramer binding in CD4^+^ and CD8^+^ cells edited with a 1G4-LY^112K^ SEED. **b,** Assessment of CD4 disruption by flow cytometry and indel generation by bulk genomic DNA sequencing in T cells after editing with intron-targeted gRNAs (*n* = 2 donors). **c**, Expression of CD4 or CD8 in subpopulations of non-purified CD8 SEED edited cells. MFI: median fluorescence intensity (*n* = 3 donors). **d,** Representative flow cytometry plots of CD4 expression, CD8 expression, BW242 binding, and NY-ESO-1 dextramer binding in cells edited with 1G4-LY SEED and CD8 SEED HDRTs. Immunomagnetically purified (anti-CD4, BW242) cells were used for longitudinal cytotoxicity assays against A375 (Fig. 4e). Colored boxes indicate subpopulations in each sample (*n* = 2 donors).

**Extended Data Figure 8:**
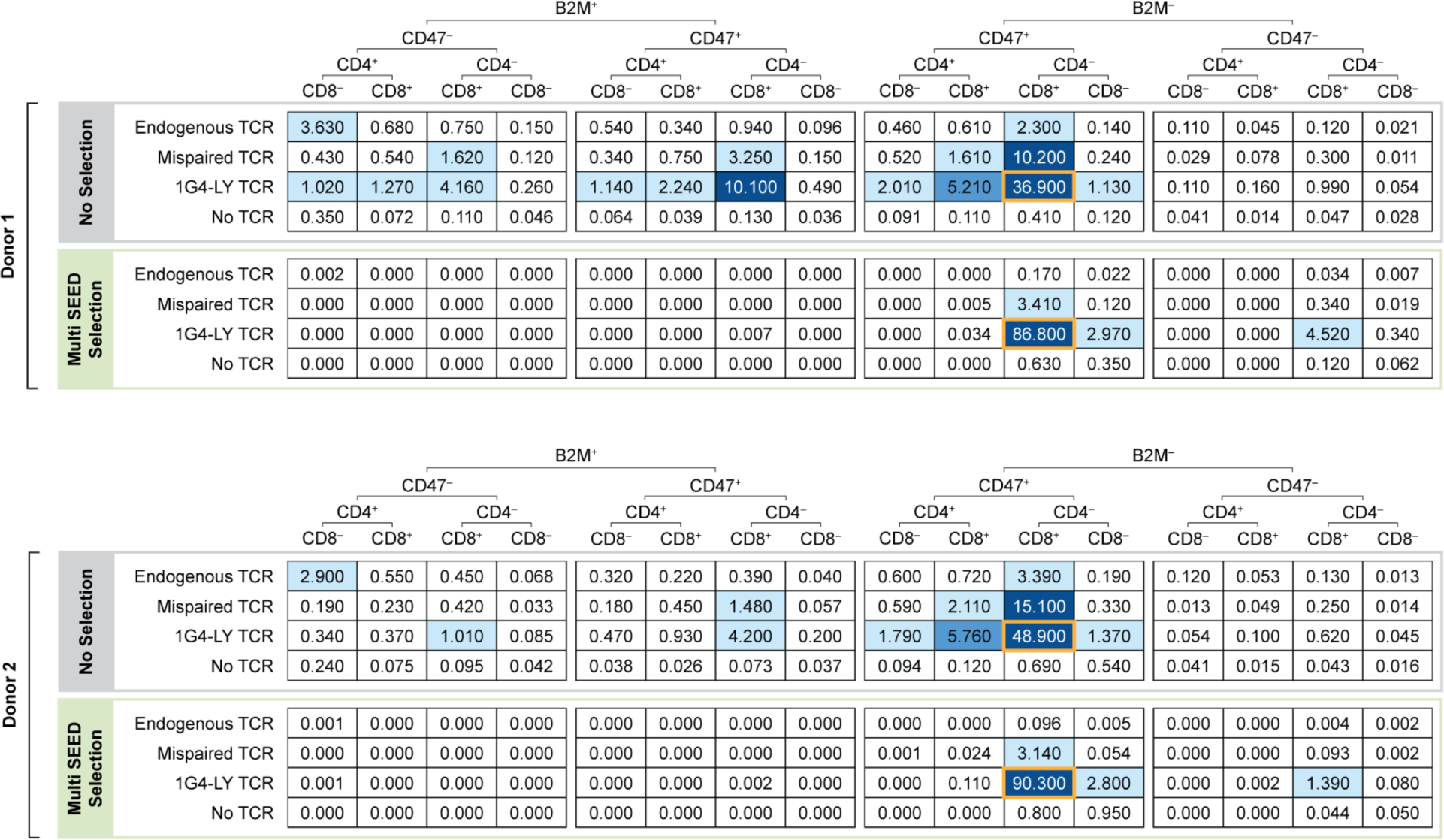
Outcomes for triple SEED engineering and SEED-selection. Outcomes after multiplexed editing with 1G4-LY^112K^, CD8, and CD47 SEED HDRTs and immunomagnetic purification with BW242, anti-CD4, and anti-B2M. Numbers indicate percentage of total population of cells. Percentage of fully-edited cells with proper 1G4-LY pairing (BW242^-^ B2M^-^ CD4^-^ Dextramer^+^ CD47^+^ CD8^+^) in each condition is enclosed by an orange box.

