## Supplementary material for "Ultra-high efficiency T cell reprogramming at multiple loci with SEED-Selection": Supplementary Information.pdf

### Contents

**Supplementary Tables:**

1. HDRT Sequences (external file)
2. Primer Sequences
3. gRNA Sequences
4. Antibodies / Dextramers Used
5. Scanning Mutagenesis Substitution Table
6. TRBC Orthologues
7. TRBC Scanning Mutagenesis Oligo Sequences (external file)

**Supplementary Figures:**

1. *TRAC*-CAR single-edit gating strategy
2. *B2M*-CD47 single-edit gating strategy
3. NK co-culture gating strategy
4. HIT$^{112K}$ gating strategy
5. Transgenic TCR gating strategy
6. *CD4*-CD8 single-edit gating strategy
7. *B2M* gDNA PCR (unedited gel image)

**Supplementary Table 2: Primer Sequences**

| **qPCR Primers** | |
|---|---|
| Sequence | Notes |
| CTTTGCTGGGCCTTTTTCCC | F primer for TRAC LHA |
| CCTGCCACTCAAGGAAACCT | R primer for TRAC LHA |
| AACATGCTACGCAGAGAGGGAGTGG | F ITR primer |
| CATGAGACAAGGAACCCCTAGTGATGGA | R ITR primer |

| **HIT Library Amplicon Sequencing Primers** | |
|---|---|
| Sequence | Notes |
| ACACTCTTTCCCTACACGACGCTCTTCCGATCTACTTTCGGTGTCAGGTCCAGTTT | F primer for HIT Amplicon + Ilumina Adapters |
| GACTGGAGTTCAGACGTGTGCTCTTCCGATCTGTGGCCTTCCCCAGGAGAAT | R primer for HIT Amplicon + Ilumina Adapters |

| **gRNA Screening Primers** | |
|---|---|
| Sequence | Notes |
| GGTGGATGAGGCACCATATTC | F primer for TRAC_i2-i10 |
| AGTCCAGATGCCAGTGATG | R primer for TRAC_i2-i10 |
| GAAGCTCAGATGCAAAGAGC | F primer for b2m_i1,i2,i3,i4 |
| CCCCTCTCTGACTTTGTACC | R primer for b2m_i1,i2,i3,i4 |
| GGTGAAATCCCGTCTCTACT | F primer for b2m_i5,i6,i7 |
| TCCACCTTCCCAACAAGCCA | R primer for b2m_i5,i6,i7 |
| GTGGCACCTGCTGAGATACT | F primer for b2m_i8,i9,i10 |
| TAATATGGCCATACCTGGGG | R primer for b2m_i8,i9,i10 |
| CTCCAAATCCAGACGCACTT | F primer for cd4_i1 |
| TCCTGGCCAGTCTCTGGTTT | R primer for cd4_i1 |
| GTAAAACGGGTTACCCAGGA | F primer for cd4_i2 |
| CCTGCACCCAGATAGTACTA | R primer for cd4_i2 |
| GCCCTGTTTCTGGTTCTGGT | F primer for cd4_i3,i6 |
| CACTCCTTAGAGGCGTATTC | R primer for cd4_i3,i6 |
| TTTGTGAGCTACTGTCCCAG | F primer for cd4_i4,i5 |
| CCCAGCCAAGATAGGGTTTC | R primer for cd4_i4,i5 |

| **HDRT Integration Site Primers** | |
|---|---|
| TCCCCAATCCACCTCTTGATG | F primer for B2M |
| GCAATTGCTCTATACGTGGCAG | R primer for B2M |

**Supplementary Table 3: gRNA Sequences**

| Target | ID | Sequence | Notes |
|---|---|---|---|
| B2M | b2m_e1 | GGCCACGGAGCGAGACATCT | |
| B2M | b2m_i1 | GCATGACTAGACCATCCATG | |
| B2M | b2m_i2 | GTGATTGCTGTAAACTAGCC | |
| B2M | b2m_i3 | TAGTTTACAGCAATCACCTG | |
| B2M | b2m_i4 | GGACCCGATAAAATACAACA | Primer selected from gRNA screen at B2M |
| B2M | b2m_i5 | CATAGCAATTGCTCTATACG | |
| B2M | b2m_i6 | TTCCTAAGTGGATCAACCCA | |
| B2M | b2m_i7 | GGAATGCTATGAGTGCTGAG | |
| B2M | b2m_i2 | GAAGCTGCCACAAAAGCTAG | |
| B2M | b2m_i9 | ACTGAACGAACATCTCAAGA | |
| B2M | b2m_i10 | ATTGTTTAGAGCTACCCAGC | |
| CD4 | cd4_i1 | GTACGTGTACGACAGTGTGT | |
| CD4 | cd4_i2 | AGCACTTGGGCTAAGAACCA | |
| CD4 | cd4_i3 | TCAGTCCTCAACTTAATACG | |
| CD4 | cd4_i4 | GGGTTTCTCTGATTAGAACG | |
| CD4 | cd4_i5 | CATCCCTCACCTGATCAAGA | |
| CD4 | cd4_i6 | TAAGTCACATAAGCACCCAG | Primer selected from gRNA screen at CD4 |
| TRAC | trac_e1 | TCAGGGTTCTGGATATCTGT | |
| TRAC | trac_i | CTGGATATCTGTGGGACAAG | |
| TRAC | trac_i2 | CAGGCACAAGCTATCAATCT | |
| TRAC | trac_i3 | AGCTATCAATCTTGGCCAAG | |
| TRAC | trac_i4 | GTGAACGTTCACTGAAATCA | |
| TRAC | trac_i5 | CTGCCAGAGTTATATTGCTG | Primer selected from gRNA screen at *TRAC* |
| TRAC | trac_i6 | AACTCTGGCAGAGTAAAGGC | |
| TRAC | trac_i7 | GTACATCTTGGAATCTGGAG | |
| TRAC | trac_i8 | CTAATGCCCAGCCTAAGTTG | |
| TRAC | trac_i9 | CTGGGCATTAGCAGAATGGG | |
| TRAC | trac_i10 | ATGGGAGGTTTATGGTATGT | |
| TRBC | | CAAACACAGCGACCTTGGGT | |

**Supplementary Table 4: Antibodies / Dextramers**

| Flow Cytometry | | | | |
|---|---|---|---|---|
| Target | Clone | Fluorophore/Marker | Manufacturer | Product # |
| TCR | BW242/412 | PE | Miltenyi | 170-081-005 |
| TCR | BW242/412 | FITC | Miltenyi | 130-113-530 |
| TCR | IP26 | BV421 | Biolegend | 306722 |
| CD4 | SK3 | BUV395 | BD | 563550 |
| CD4 | SK3 | FITC | BD | 344604 |
| CD8 | SK1 | BV421 | BD | 740093 |
| CD8 | SK1 | PE-Cy7 | BD | 335787 |
| G4S Linker | E7O2V | PE | Cell Signaling | 38907L |
| Mouse F(ab')2 | N/A | AF647 | Jackson ImmunoResearch | 115-606-072 |
| B2M | 2M2 | APC | Biolegend | 316311 |
| CD47 | B6H12 | PerCP-Cy5.5 | BD | 561261 |
| CD19 | SJ25C1 | BUV373 | BD | 612756 |
| CD56 | My31.13 | FITC | UCSF Monoclonal Antibody Core | |
| CD3 | UCHT1 | BV711 | Biolegend | 300464 |
| NY-ESO-1 Dextramer (SLLMWITQV) | N/A | PE | Immudex | WB03247 |
| MART-1 Dextramer (ELAGIGILTV) | N/A | PE | Immudex | WB02162 |

| Negative Selection | | | | |
|---|---|---|---|---|
| Target | Clone | Fluorophore/Marker | Manufacturer | Product # |
| TCR | BW242/412 | Biotin | Miltenyi | 130-021-301 |
| B2M | 2M2 | Biotin | Biolegend | 316308 |
| CD4 | SK3 | Biotin | Biolegend | 344610 |

**Supplementary Table 5: Scanning Mutagenesis Substitution Table**

| Original AA | A | R | N | D | C | Q | E | G | H | I | L | K | M | F | P | S | T | W | Y | V |
|---|---|---|---|---|---|---|---|---|---|---|---|---|---|---|---|---|---|---|---|---|
| **Substitutions** | S | K | D | N | T | E | Q | N | Y | V | M | R | L | Y | K | T | S | Y | F | I |
| | G | Q | Q | E | V | R | D | S | Q | L | I | Q | I | L | S | N | V | F | H | L |
| | T | H | H | Q | I | H | K | D | R | M | V | E | V | M | R | Q | N | L | W | M |
| | C | N | K | K | L | K | H | Q | N | F | F | N | Q | W | D | E | R | M | Q | T |
| | V | A | A | A | A | A | A | A | A | A | A | A | A | A | A | A | A | A | A | A |

**Supplementary Table 6: TRBC Orthologues**

1. 0A5H1ZRT1_HUMAN
2. 0A2J8QUE2_PANTR
3. 2PP94_PONAB
4. 0A2J8V300_PONAB
5. 0A0G2JMB4_HUMAN
6. 0A5H1ZRR3_HUMAN
7. 0A0D9R5B1_CHLSB
8. 0A2I3SCJ3_PANTR
9. 7MMZ5_MACMU
10. 0A2R8ZDC5_PANPA
11. 0A2I2YRH4_GORGO
12. 0A5E4AQB3_MARMO
13. 1PSA6_MYOLU
14. 0A4W2ERJ9_BOBOX
15. 1MJJ6_BOVIN
16. 1MUR8_BOVIN
17. 3MXJ1_BOVIN
18. 8IAU9_9CETA
19. 0A452E0J8_CAPHI
20. 0A452E0R0_CAPHI
21. 1PEQ3_CANLF
22. 0A452ERL6_CAPHI
23. 0A2K6GVW3_PROCO
24. 0A2K6GVV9_PROCO
25. 2HP35_AILME
26. 1LIB9_AILME
27. 2I8R3_AILME
28. 2HP36_AILME
29. 0A4W2HJF4_BOBOX
30. 1MJB8_BOVIN
31. 0A673SS53_SURSU
32. 0A673SS69_SURSU
33. 0A485NS05_LYNPA
34. 3YAT2_MUSPF
35. 0A3P4NJG5_GULGU
36. 0A2K5PVI3_CEBCA
37. 0A2K5PVI6_CEBCA
38. 0A2K6TGF3_SAIBB

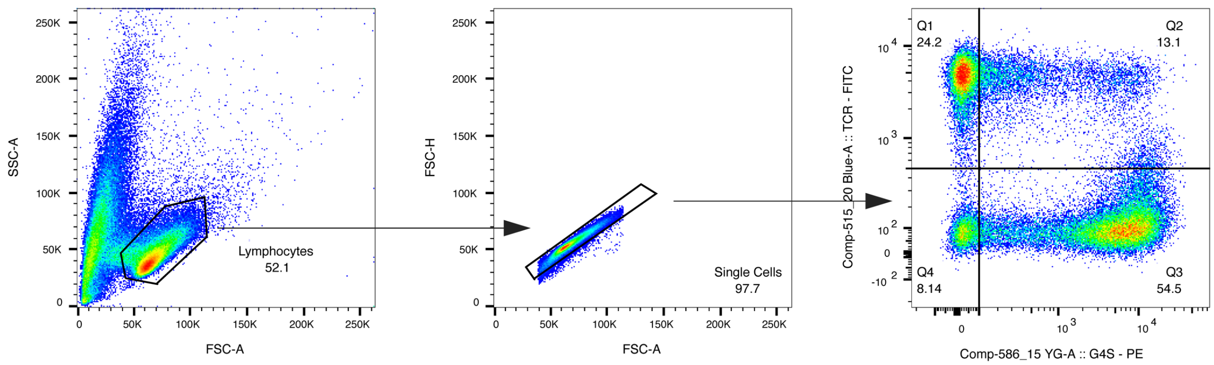


**Supplementary Figure 1:** General gating strategy for assessing editing outcomes in T cells edited with *TRAC*-targeted HDRTs encoding CARs. Gates for CAR and TCR expression were set based on edited non-transduced controls and non-edited controls.

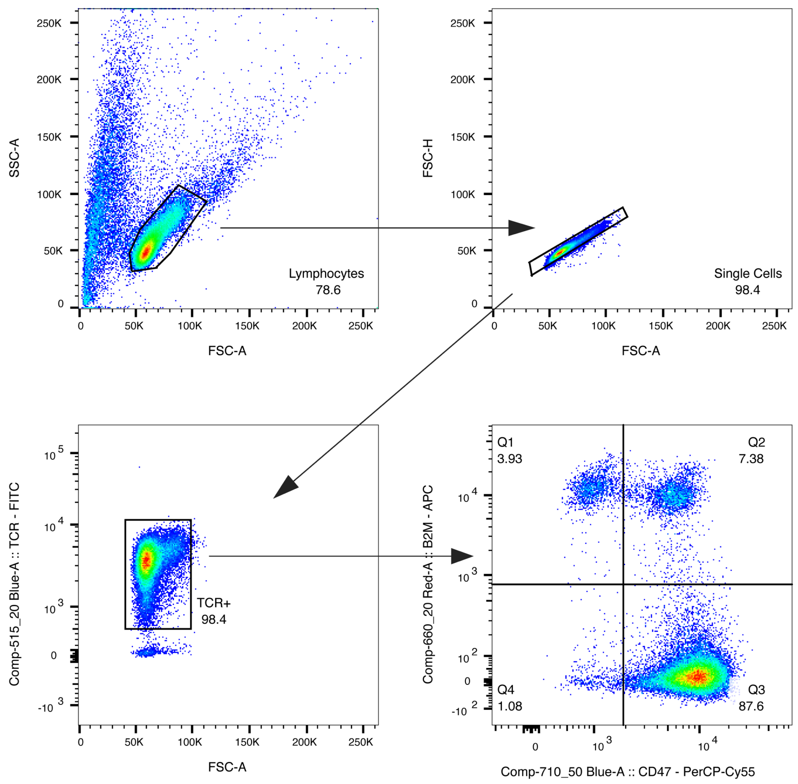


**Supplementary Figure 2:** Gating strategy for assessing editing outcomes in T cells edited with *B2M*-targeted HDRTs encoding CD47. Gates for B2M and CD47 expression were set based on edited non-transduced controls and unedited controls.

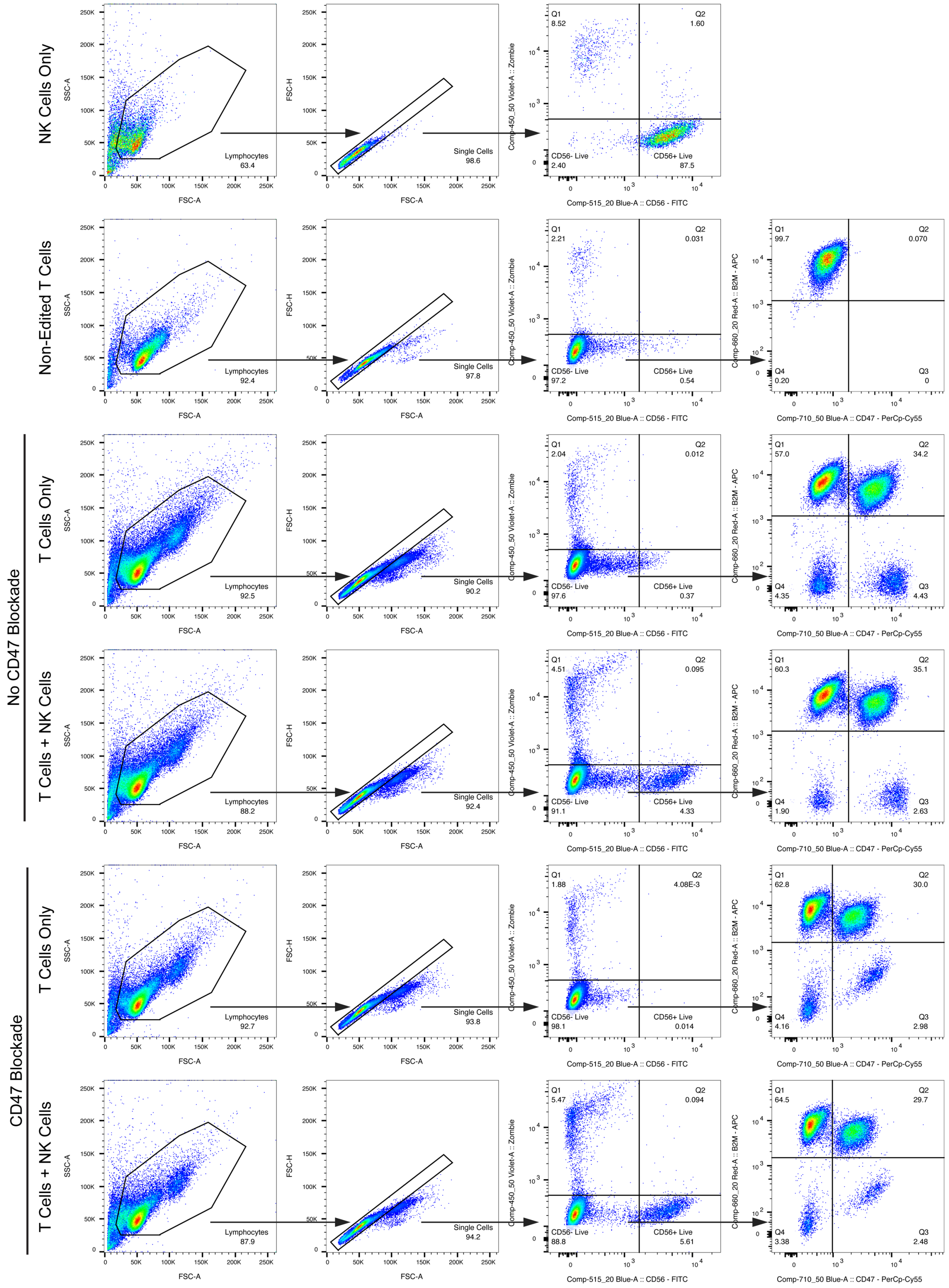


**Supplementary Figure 3:** Gating strategy for NK cell co-cultures. Gating for CD56 was set based on an NK cell only control sample. Separate gates for B2M and CD47 expression were used for samples treated with anti-CD47 and non-treated samples.

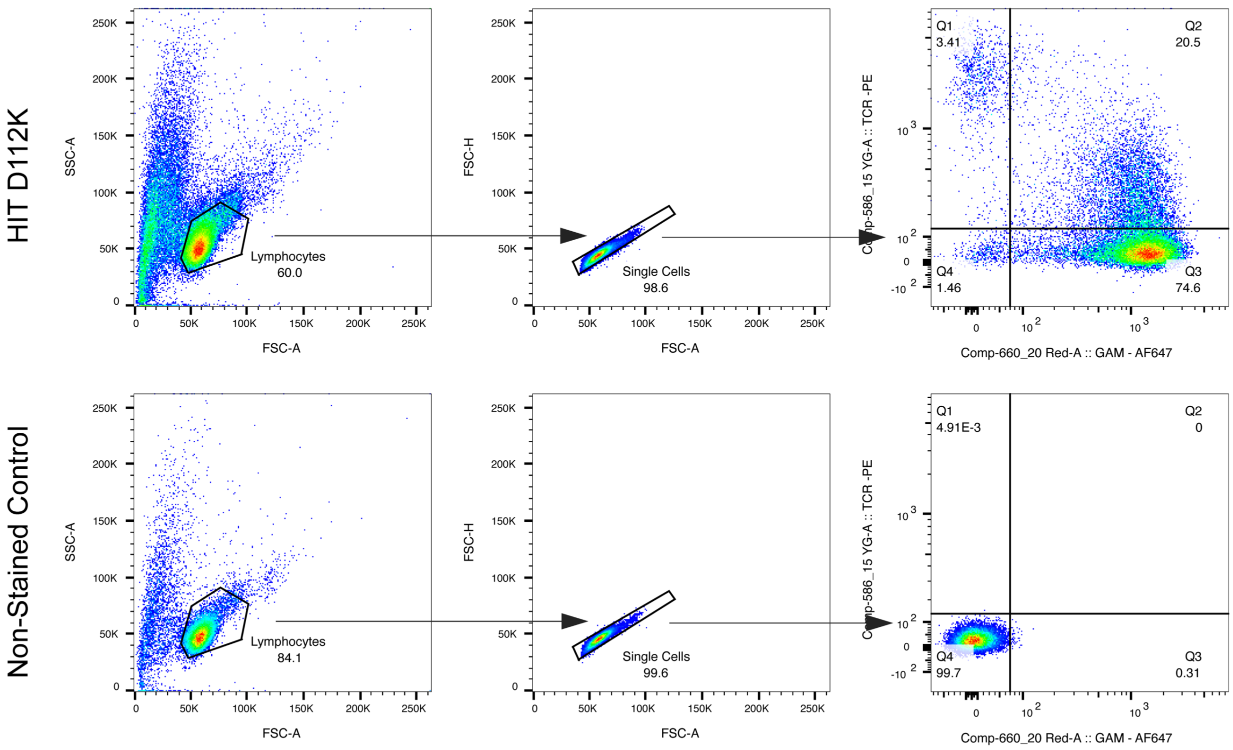


**Supplementary Figure 4:** Gating strategy for assessing editing outcomes in T cells edited with a *TRAC*-targeted HDRT encoding HIT D112K. Gating for TCR and HIT expression was set based on edited non-transduced samples and non-edited samples.

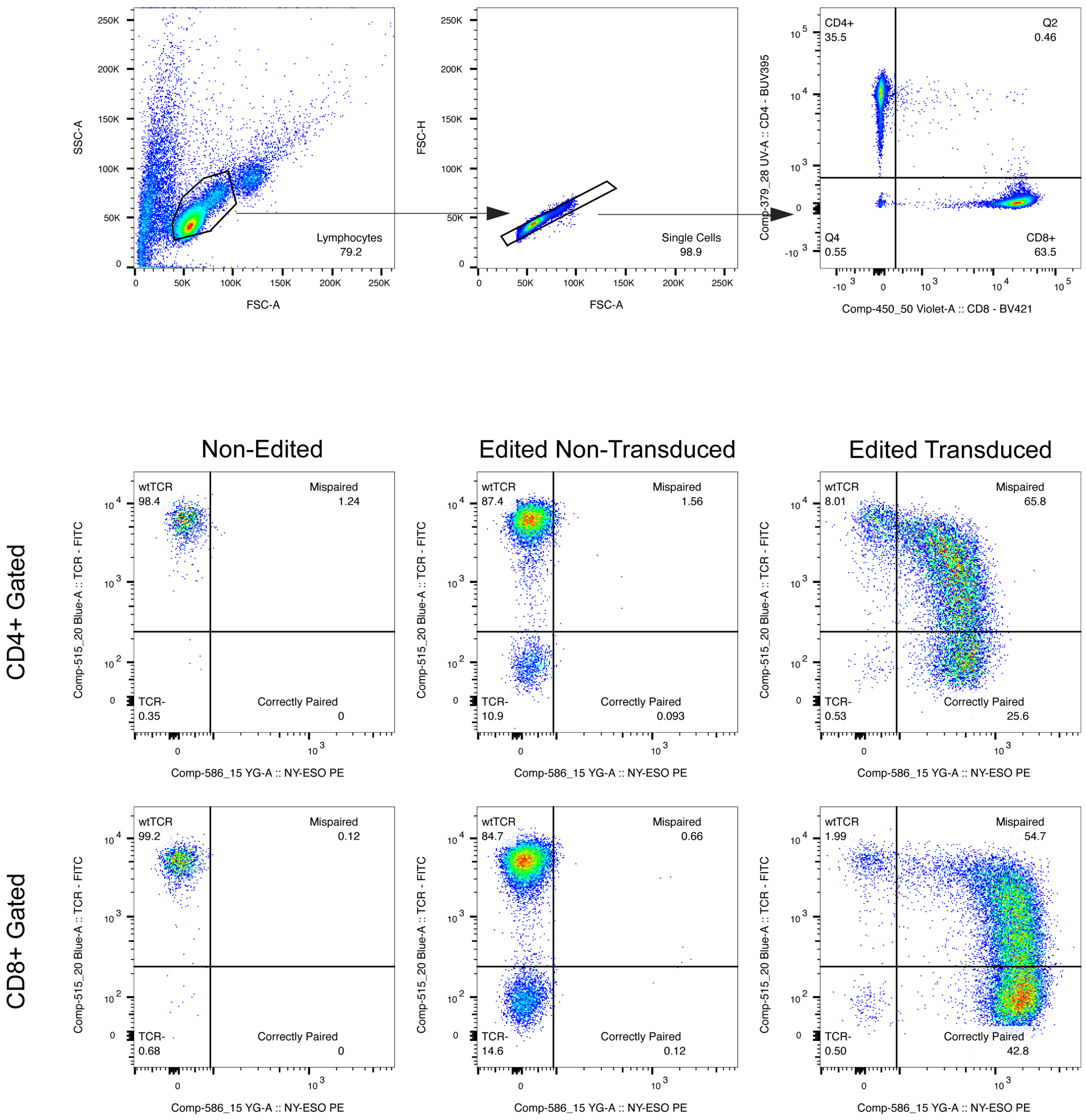


**Supplementary Figure 5:** General gating strategy for assessing editing outcomes and mispairing in T cells edited with a *TRAC*-targeted HDRT encoding an epitope edited transgenic TCR. Gating for mispairing were set based on edited non-transduced control samples.

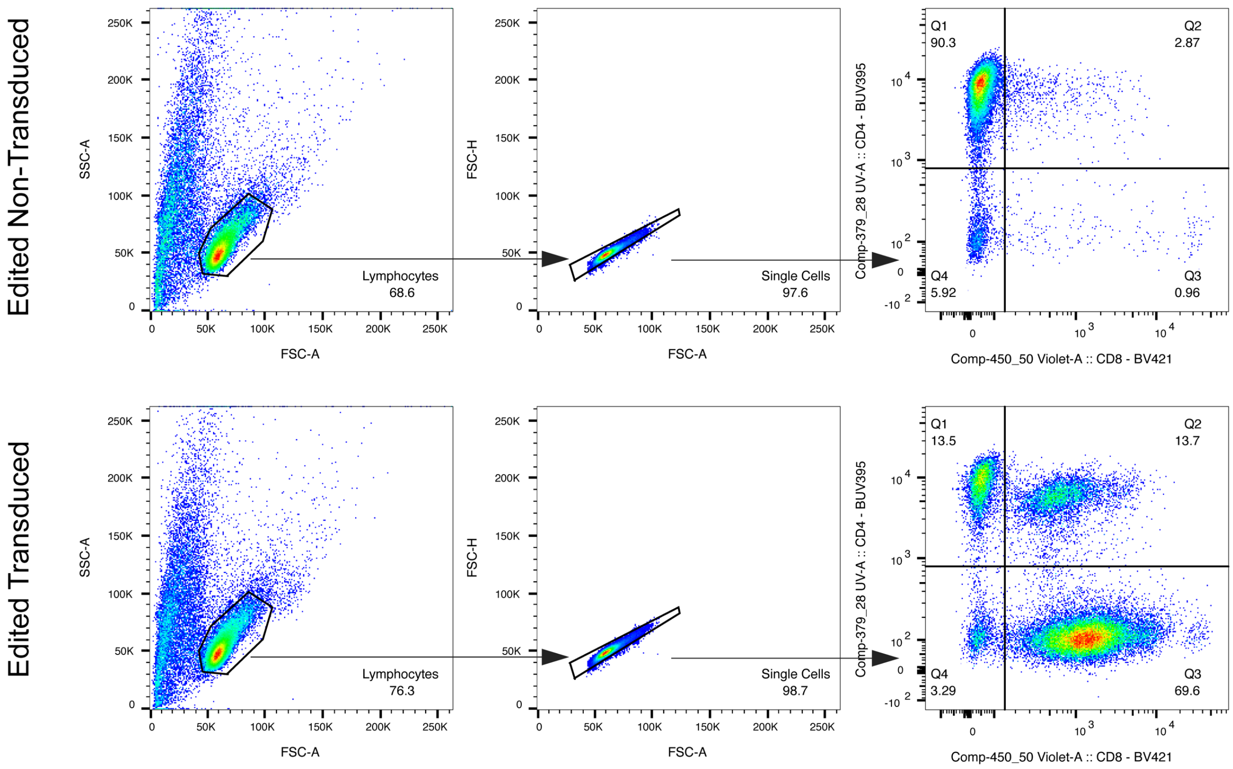


**Supplementary Figure 6:** Gating strategy for assessing editing outcomes in CD4⁺ T cells edited with a *CD4*-targeted HDRT encoding CD8. Gating for CD4 and CD8 expression was set based on edited non-transduced controls.

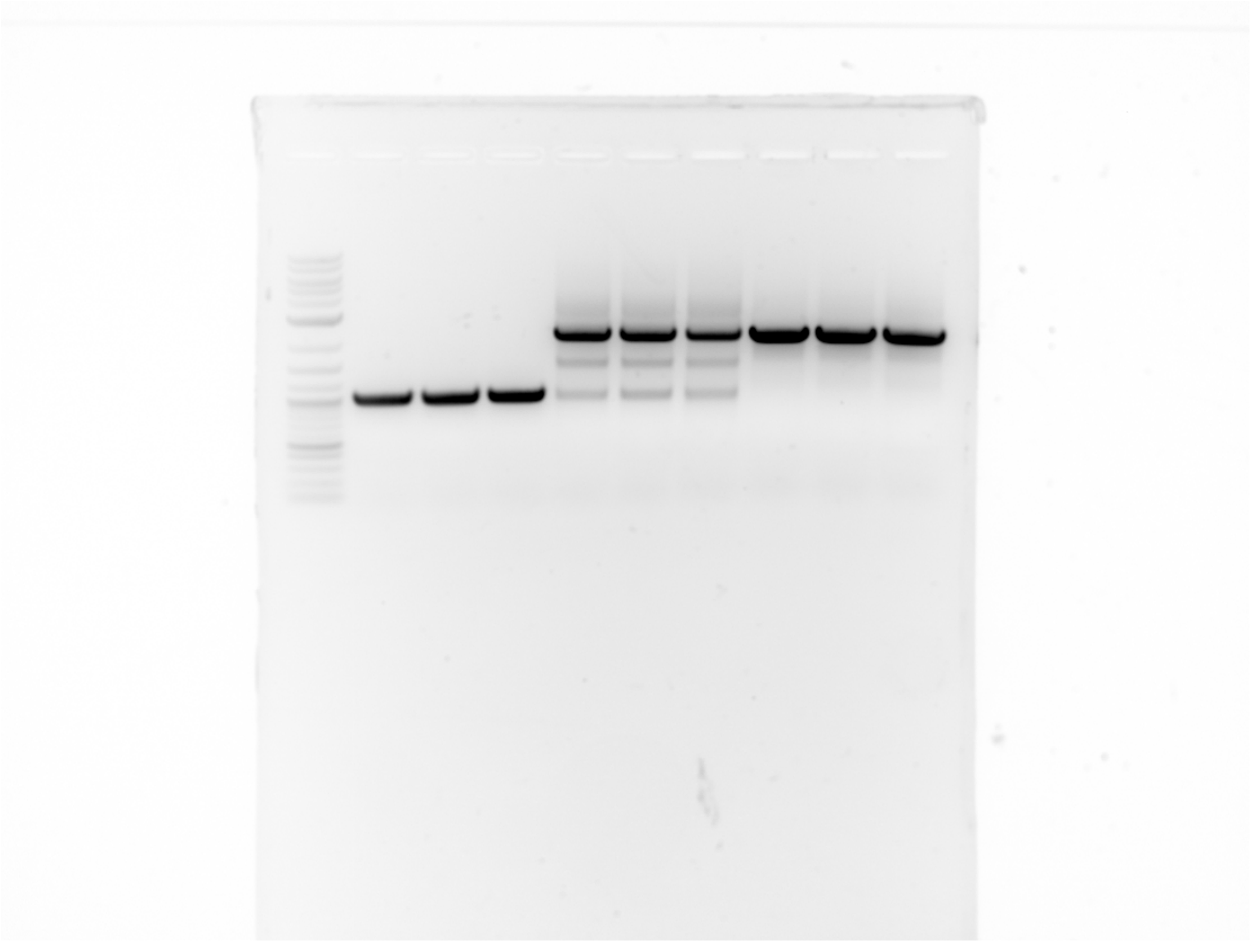

**Supplementary Figure 7:** Unprocessed gel image of *B2M* HDRT integration site gDNA PCR (Cropped gel shown in Fig. 1i).
